# Structural and mechanistic basis for protein glutamylation by the kinase fold

**DOI:** 10.1101/2021.04.13.439722

**Authors:** Adam Osinski, Miles H. Black, Krzysztof Pawłowski, Zhe Chen, Yang Li, Vincent S. Tagliabracci

## Abstract

The kinase domain transfers phosphate from ATP to substrates. However, the *Legionella* effector SidJ adopts a kinase fold yet catalyzes calmodulin (CaM)-dependent glutamylation to inactivate the SidE ubiquitin ligases. The structural and mechanistic basis in which the kinase domain catalyzes protein glutamylation is unknown. Here we present cryo-EM reconstructions of SidJ:CaM:SidE reaction intermediate complexes. We show that the kinase-like active site of SidJ adenylates an active site Glu in SidE resulting in the formation of a stable reaction intermediate complex. An insertion in the catalytic loop of the kinase domain positions the donor Glu near the acyl-adenylate for peptide bond formation. Our structural analysis led us to discover that the SidJ paralog SdjA is a glutamylase that differentially regulates the SidE-ligases during *Legionella* infection. Our results uncover the structural and mechanistic basis in which the kinase fold catalyzes non-ribosomal amino acid ligations and reveal an unappreciated level of SidE-family regulation.

## Introduction

*Legionella pneumophila*, the causative agent of Legionnaires disease, translocates more than 330 effectors utilizing a Type 4 secretion system to establish a replicative niche known as the *Legionella* Containing Vacuole (LCV) (Cornejo et al., 2017; Isberg et al., 2009). Within the *Legionella* effector repertoire are protein domains with recognizable folds that occasionally catalyze unexpected reactions (Black et al., 2019; Mukherjee et al., 2011; Neunuebel et al., 2011; Qiu et al., 2016). For example, the *Legionella* effector SidJ adopts a kinase-like fold yet catalyzes calmodulin (CaM)-dependent glutamylation to inactivate the SidE ubiquitin (Ub) ligases (Bhogaraju et al., 2019; Black et al., 2019; Gan et al., 2019; Sulpizio et al., 2019). The SidE effectors SdeA, SdeB, SdeC and SidE (collectively referred to as SidE), employ ADP ribosyltransferase (ART) and phosphodiesterase (PDE) activities to catalyze ligation of Ub to proteins independent of E1 and E2 enzymes (Bhogaraju et al., 2016; Kotewicz et al., 2017; Qiu et al., 2016). The SidE ART domain uses NAD+ to ADP-ribosylate Ub on Arg42. ADP ribosylated Ub serves as a substrate for the PDE domain, which hydrolyzes the phosphodiester bond and transfers Ub to Ser residues on proteins forming a Ser-phosphoribosyl (pR)-Ub linkage (Akturk et al., 2018; Dong et al., 2018; Kalayil et al., 2018; Wang et al., 2018).

SidE-family ubiquitination promotes infectivity of *Legionella pneumophila* and is required for proper formation of the LCV (Bardill et al., 2005). However, unrestrained SidE activity is harmful to the host. Therefore, SidE-family ubiquitination is regulated by *Legionella* deubiquitinases DupA and DupB, which reverse pR ubiquitination (Shin et al., 2020; Wan et al., 2019), and the pseudokinase SidJ, which glutamylates and inactivates the SidE effectors (Bhogaraju et al., 2019; Black et al., 2019; Gan et al., 2019; Sulpizio et al., 2019). Three of the four SidE-family members lie within a contiguous genomic locus that also includes *dupA* and *sidJ* (**Figure 1A**). The *dupB* gene neighbors the *sidJ* paralog, *sdjA*. Despite sharing 52% amino acid sequence identity, SidJ and SdjA are not functionally redundant (Qiu et al., 2017b). As such, the function of SdjA is unknown.

**Figure 1.**
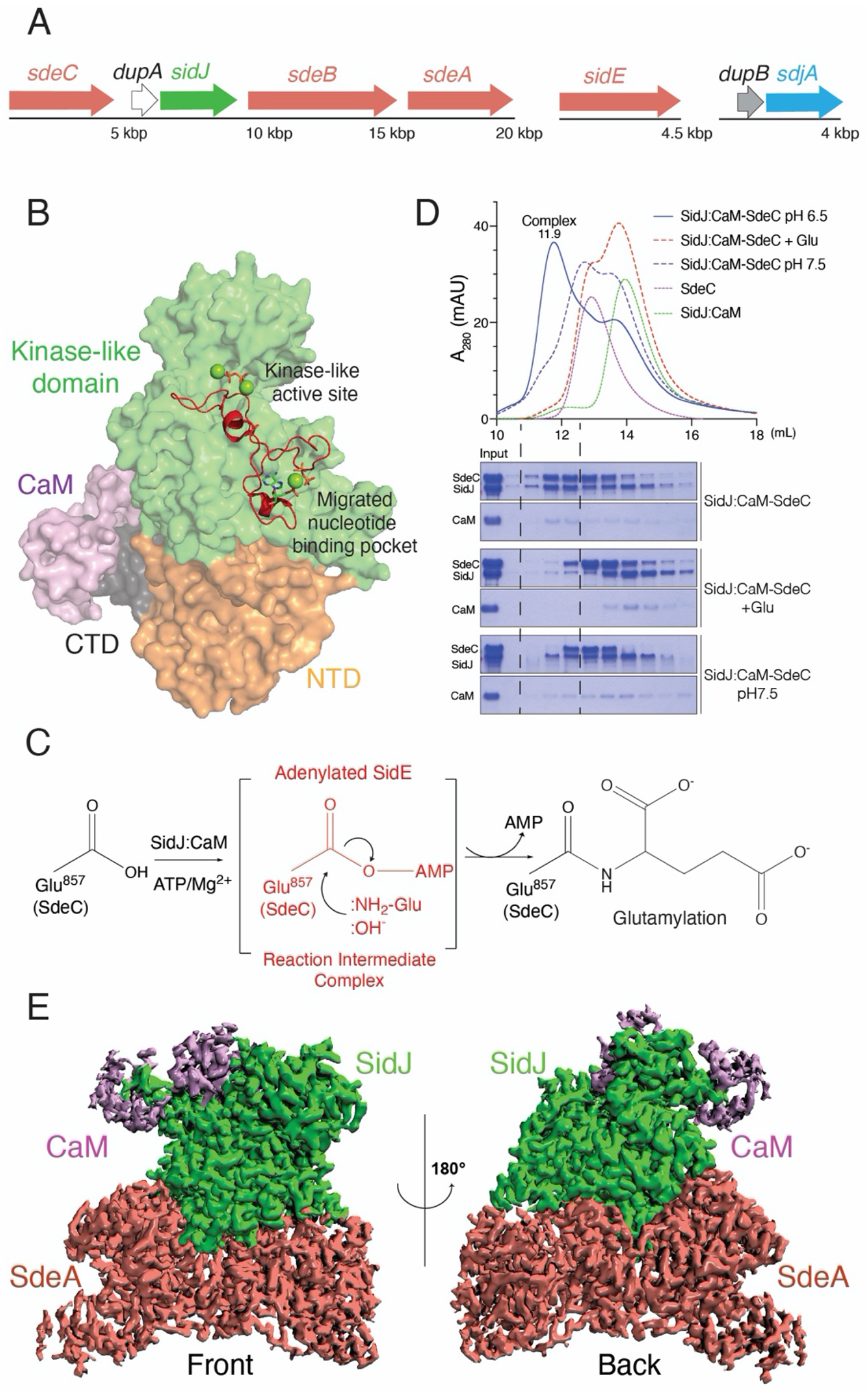
SidJ forms a stable acyl-adenylate-intermediate complex with SdeA and SdeC. **(A)** Organization of the *sidE* family (salmon), *sidJ* (green) and *dupA* (white), *dupB*, (grey) and *sdjA* (blue) effectors in the genome of *L. pneumophila*. SdeA, SdeB and SdeC lie in a genomic neighbourhood with SidJ and DupA, while SidE lies in a different locus (upper). **(B)** Structure of SidJ depicting the N-terminal domain (NTD; orange), the kinase-like domain (green), the C-terminal domain (CTD; black) and CaM (purple). The kinase-like active site and the migrated nucleotide binding pocket formed by an insertion in the catalytic loop (red) are highlighted. **(C)** Schematic representation of SidJ-catalyzed glutamylation of the SidE effectors. The SidJ:CaM complex binds ATP/Mg^2+^, which is used to adenylate the SidE-family active site Glu (SdeC; Glu^857^). Adenylated SidE forms a stable reaction intermediate complex with SidJ, which facilitates Glu binding, positioning of the NH_2_ group for attack of the acyl-adenylate and formation of the isopeptide bond. Note that under basic conditions, :OH^−^ can hydrolyze the acyl-adenylate. **(D)** SEC trace (upper) and SDS-PAGE and Coomassie staining (lower) of the SidJ-SdeC complex (blue) and the products resulting from the addition of Glu (dashed red) or increasing the pH to 7.5 (dashed blue). Note that the addition of Glu and increasing the pH dissociates the complex, consistent with the presence of an acyl-adenylate in the complex. **(E)** Cryo-EM density map representation of the SidJ:CaM:SdeA complex. SidJ is in green, SdeA^Core^ is in salmon and CaM is in purple. Front and back views are shown.

Pseudoenzymes contain a protein domain that resembles a catalytically active counterpart (Kwon et al., 2019; Ribeiro et al., 2019). However, pseudoenzymes lack key catalytic residues believed to be required for activity. Recent work on pseudokinases has revealed that different binding orientations of ATP and active site residue migration can repurpose the kinase scaffold to catalyse novel reactions (Black et al., 2019; Sreelatha et al., 2018). These results suggest that the kinase fold is more versatile than previously appreciated and that pseudokinases should be reanalyzed for alternative transferase activities.

Structures of the SidJ:CaM complex uncovered an N-terminal domain (NTD) of unknown function, a kinase-like domain with two ligand binding pockets and a C-terminal domain that binds CaM (Bhogaraju et al., 2019; Black et al., 2019; Gan et al., 2019; Sulpizio et al., 2019) (**Figure 1B**). The canonical kinase-like ATP binding pocket of SidJ contains residues that are required for glutamylation of the SidE ligases. An insertion in the catalytic loop of the kinase domain extends away from the kinase active site and forms a migrated nucleotide binding pocket, which is also required for glutamylation (Bhogaraju et al., 2019; Black et al., 2019; Gan et al., 2019; Sulpizio et al., 2019). From these studies, we proposed a catalytic mechanism, whereby CaM binding allows the kinase–like active site of SidJ to bind ATP and transfer AMP to the active site Glu on SidE forming a high energy acyl-adenylate intermediate. Adenylated SidE binds the migrated nucleotide-binding pocket, positioning the acyl-adenylate and donor Glu for glutamylation and inactivation of the SidE effectors (Black et al., 2019) (**Figure 1C**). However, others argue that the migrated nucleotide binding pocket is an allosteric site that facilitates glutamylation (Sulpizio et al., 2019).

Here, we present cryo-EM structures of the reaction intermediate complex formed between SidJ and SdeA, and SidJ and SdeC. We propose a model that explains how a pseudokinase catalyzes protein glutamylation. Furthermore, we show that the SidJ paralog, SdjA is an active glutamylase that differentially regulates SidE-family activity in vitro and during *Legionella* infection.

## Results

### SidJ and SdeA/C Form a Stable Reaction Intermediate Complex

In several amidoligases, including the ATP-grasp enzymes involved in tubulin glutamylation, a stable reaction intermediate complex facilitates the formation of the peptide bond (Garnham et al., 2015; Mahalingan et al., 2020). We hypothesized that SidJ:CaM would also form a stable reaction intermediate complex with the SidE ligases when Glu was excluded and when the reaction was performed under acidic conditions to prevent base-catalyzed hydrolysis of the acyl-adenylate (**Figure 1C**). We incubated SidJ^97-851^:CaM (SidJ:CaM) with SdeA^231-1190^ (SdeA^Core^) or SdeC^231-1222^ (SdeC^Core^), initiated the reactions with ATP/Mg^2+^, and analyzed the reaction products by SDS-PAGE following size-exclusion chromatography (SEC) in a slightly acidic (pH 6.5) buffer. We detected the formation of stable ternary complexes consisting of SidJ, CaM, and SdeC^Core^ (**Figure 1D**) and SidJ, CaM and SdeA^Core^ (**Figure S1A**). Upon supplementation of the reaction with Glu, or when the complex was subjected to SEC using a mildly basic buffer (pH 7.5), the SidJ:CaM:SdeC^Core^ complex dissociated (**Figure 1D**). These results are consistent with the presence of an acyl-adenylate intermediate within the complex.

### Extensive Interface Interactions Facilitate Complex Formation Between SidJ:CaM and SdeA/C^Core^

We determined cryo-EM structures of SidJ:CaM:SdeA^Core^ and SidJ:CaM:SdeC^Core^ complexes with resolution up to 2.5 Å and 2.8 Å, respectively (**Figure 1E, and Figures S1-S4, Table S1**). The density maps reveal details of the interface between the proteins and their active sites, with clear densities for SidJ, CaM and the PDE and ART domains of SdeA^Core^ and SdeC^Core^. Densities for the distal parts of the PDE domain and most of the C-terminal coiled coil are less clear, suggesting a higher degree of flexibility in these regions. We use the SidJ:CaM:SdeA^Core^ maps for description of the interface and the SidJ:CaM:SdeC^Core^ maps to describe interactions related to catalysis.

The overall reaction intermediate complexes are highly similar, with a root mean square deviation (RMSD) of 1.5 Å (**Figures S4C and S5**). The SidJ/CaM conformation remains similar to those previously determined by X-ray crystallography (Black et al., 2019) and cryo-EM (Bhogaraju et al., 2019). SidJ sits in a V-shaped cleft made up of the PDE and ART domains of SdeA/C^Core^ (**Figure 2A**). The interfaces between SidJ and SdeA/C^Core^ span ~2,200 Å^2^ and are predominantly formed by the SidJ NTD (**Figure 2B**). Within the NTD of SidJ, there are three regions that form extensive contacts with SdeA^Core^. **1)**A helix-turn-helix (HTH) motif that forms electrostatic contacts with residues located in the back of the ART domain, **2)**a *β*-hairpin wedged in between the PDE and ART domains and **3)**a helical bundle that interacts with the PDE domain. To verify the relevance of these interactions for SidJ-catalyzed glutamylation of SdeA, we mutated D833, K569, D565 and D372 of SdeA^Core^, which lie near the interfaces of the three regions. Reversing the charge on these residues in SdeA^Core^ markedly reduced its propensity to be glutamylated by SidJ (**Figure 2C**). Our observation that SidJ interacts with both the SdeA/C ART and PDE domains reconciles previous data showing that SidJ-dependent suppression of SdeA-mediated yeast toxicity requires the PDE and ART domains of SdeA (Havey and Roy, 2015). As expected, SidJ did not glutamylate the isolated ART domain of SdeA (**Figure 2D**).

**Figure 2.**
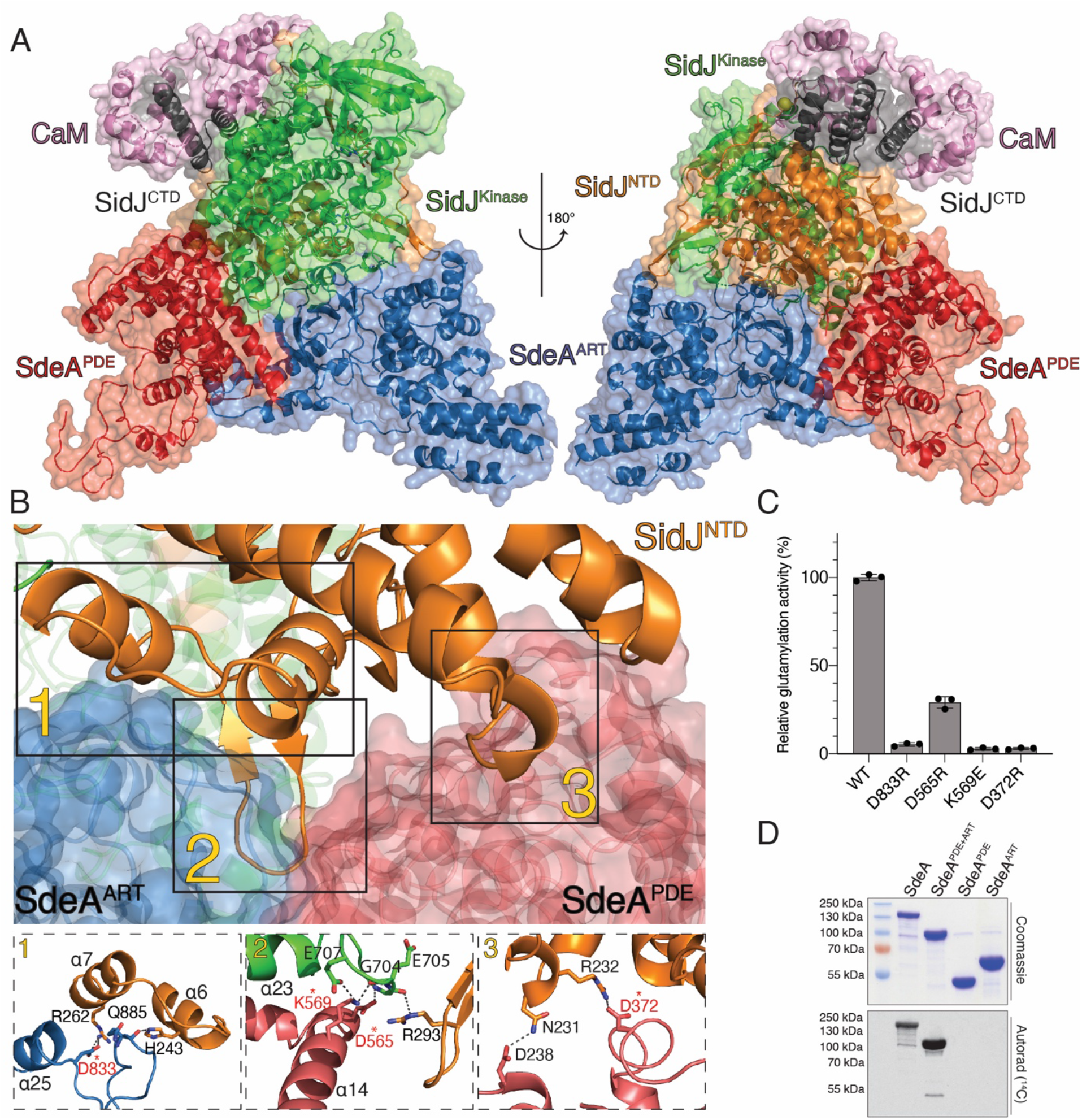
SidJ:CaM:SdeA/C^Core^ complex formation is mediated by extensive interface interactions. **(A)** Overview of SidJ:CaM:SdeA^Core^ reaction intermediate complex. The SidJ NTD, kinase-like domain and CTD are in orange, green, and black, respectively. The SdeA PDE domain is in red and the ART domain is in blue. CaM is in purple. SidJ lies within a V-shaped cleft between SdeA PDE and ART domains. **(B)** Back view of the SidJ:CaM:SdeA^Core^ interface. The SidJ NTD forms three distinct regions of interaction with SdeA (labelled 1-3). **1.** A loop within the SdeA ART domain interacts with the helix-turn-helix (HTH) motif within the SidJ NTD. **2.** A β-hairpin from the SidJ NTD is nestled between the PDE and ART domains of SdeA, forming electrostatic and hydrophobic interactions. A helix from the SidJ kinase-like domain also interacts with the SidJ β-hairpin and the PDE domain of SdeA. **3.** A helical bundle from the SidJ NTD interacts with the PDE domain of SdeA. Zoomed in view depicting the residues involved in the interaction interfaces are shown in subpanels (lower) and colored as in **A.** Residues targeted for mutagenesis are marked with asterisks. Putative hydrogen bonds and salt bridges are depicted as dotted lines. **(C)** Glutamylation activity of SidJ using SdeA^Core^ and mutations that disrupt interface interactions. Residues mutated are also marked with an asterisk in **(B)**. [^3^H]Glu was used as a substrate and the reaction products were resolved by SDS PAGE and visualized by Coomassie staining. Radioactive gel bands were excised and ^3^H incorporation into SdeA^Core^ was quantified by scintillation counting. **(D)** Incorporation of [^14^C]-Glu into full length SdeA, SdeA^PDE+ART^, SdeA^PDE^ or SdeA^ART^ by SidJ. Reaction products were separated by SDS PAGE and visualized by Coomassie staining (upper) and autoradiography (lower).

### Unique Active Sites in SidJ Catalyze Adenylation and Glutamylation

Within the kinase-like domain, the α-23 helix from the C-lobe of SidJ interacts with the α-14 helix within the PDE domain of SdeA^Core^, adjacent to its interaction with the NTD β-hairpin (**Figure 2B; inset 2**). Interactions also occur within the migrated-nucleotide binding pocket and the ARTT loop within the active site of the ART domain. Remarkably, we observed density that is consistent with a covalent bond between the phosphate of AMP and the γ-carboxyl group of E857 in SdeC^Core^, which is consistent with an acyl-adenylate (**Figure 3A)**. The acyl-adenylate is stabilized by R500, N733, H492 and Y506 within the SidJ migrated nucleotide binding pocket. Mutation of these residues to Ala markedly reduced glutamylation of SdeA by SidJ (Black et al., 2019). When comparing the active site of SdeA in the reaction intermediate complex to the apo structure of SdeA (Dong et al., 2018), significant rearrangement of the ARTT loop is observed, which facilitates SidJ binding and stabilization of the acyl-adenylate (**Figure S6A**).

**Figure 3.**
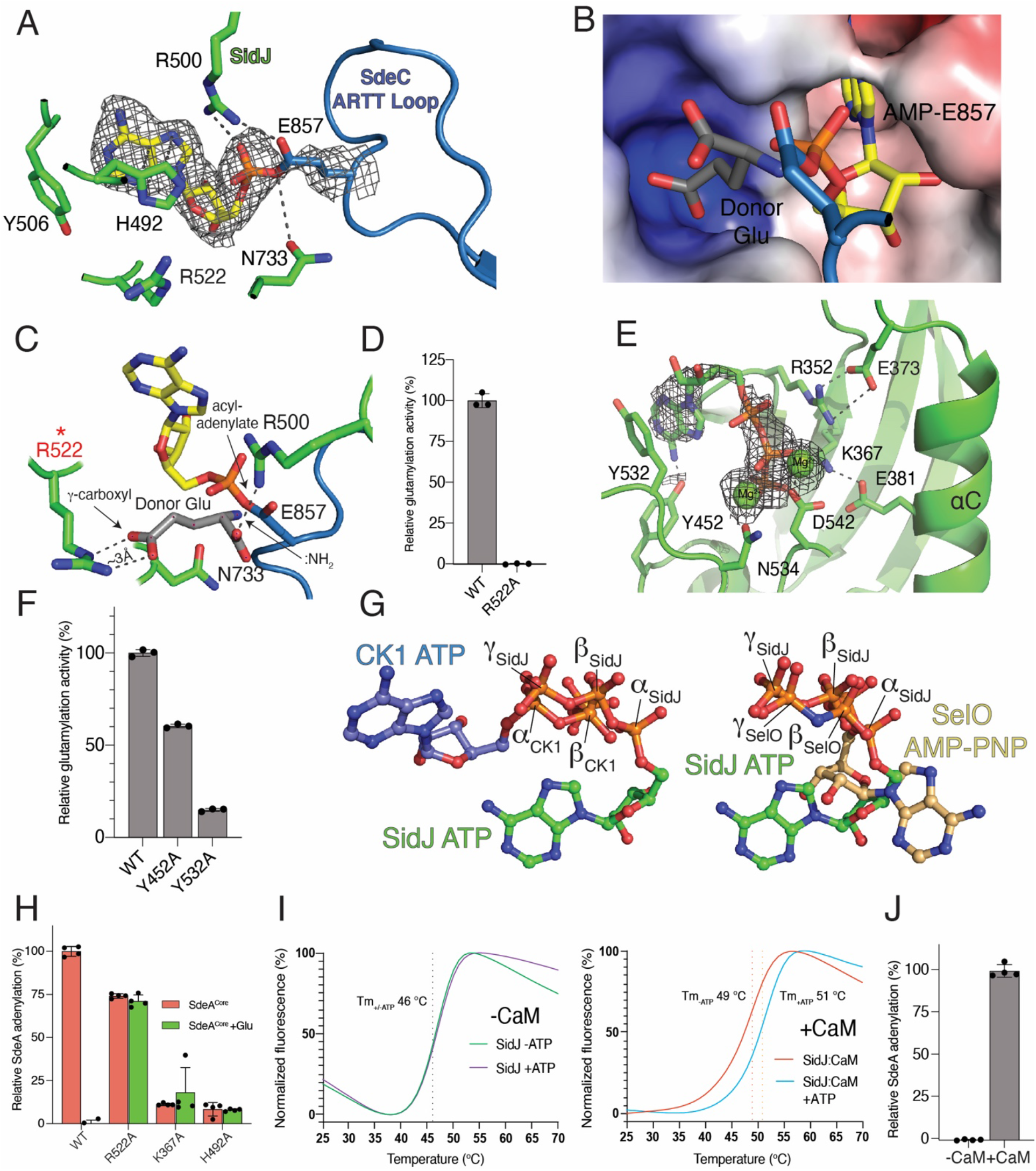
Unique active sites facilitate SidJ-mediated glutamylation of the SidE effectors. **(A)** Details of the migrated nucleotide binding pocket of SidJ (green) and the ARTT loop of SdeC^Core^ (blue) showing the interactions (dashed lines) involved in binding the acyl-adenylate reaction intermediate. The AMP is shown as sticks and the Coulomb potential map is shown in mesh. Note the formation of a covalent bond between AMP and the SdeC active site Glu857. **(B, C)** Electrostatic surface representation **(B)** and molecular interactions **(C)** of the migrated nucleotide binding pocket of SidJ bound to SdeC^Core^ depicting a positively charged cleft adjacent to the acyl-adenylate reaction intermediate. A donor Glu was manually placed inside of the pocket, with its γ-carboxyl group interacting with Arg522 of SidJ. The donor Glu :NH_2_ group is in position to attack the acyl-adenylate. Electrostatic surface is contoured at 7kT. **(D)** Glutamylation activity of SidJ and the Arg522 mutant using SdeA^Core^ as substrate. Reaction products were analyzed as in **Figure 2C**. **(E)** Details of the kinase-like active site of SidJ (green) depicting the molecular interactions (dashed lines) that facilitate ATP/Mg^2+^ binding. The ATP is in sticks, the Coulomb potential map is shown in mesh and the Mg^2+^ ions are shown as spheres. **(F)** Glutamylation activity of SidJ and kinase-like active site mutants using SdeA^Core^ as a substrate. Reaction products were analyzed as in **Figure 2C**. Note that the other residues have been mutated and analyzed for glutamylation in our previous work (Black et al., 2019). **(G)** Ball-and-stick representation of superimposed nucleotides from SidJ and protein kinase CK1 (left) and SidJ and SelO (right) as a result of superposition of the kinase active sites. The α, β, and γ-phosphates of SidJ, SelO and CK1 are highlighted. Note that the nucleotide orientation of SidJ is similar to SelO, which transfers AMP to proteins (Sreelatha et al., 2018). **(H)** Quantification of acyl-adenylate formation following reactions with SidJ^59-851^ (WT and mutants), SdeA^Core^ and [α-^32^P]ATP. SdeA^Core^ E860Q was used to calculate the baseline signal in TCA precipitates. The reactions were terminated by the addition of TCA and the SdeA acyl-adenylate was detected by scintillation counting of the acid-insoluble material (red bars). Glu was also added to the reactions (green bars), which displaces the ^32^P-AMP from SdeA into a TCA soluble fraction. **(I)** Differential scanning fluorimetry (DSF) depicting the thermal stability profiles of SidJ (left) and SidJ:CaM (right) in the presence or absence of ATP/Mg^2+^. Protein denaturation was followed by monitoring the fluorescence of SYPRO Orange dye, which binds hydrophobic regions on proteins. The T_m_ values are shown in the insets. **(J)** Quantification of acyl-adenylate formation following reactions with SidJ^59-851^, SdeA^Core^, [α-^32^P]ATP in the presence (+) or absence (−) of CaM. SdeA^Core^ E860Q was used to calculate the baseline signal in TCA precipitates. Reaction products were analyzed as in **Figure 3H**.

During catalysis, SidJ cannot substitute Asp for Glu, suggesting a high degree of specificity (Black et al., 2019). Analysis of the electrostatic surface in the vicinity of the active site revealed a small positively charged pocket in SidJ that we predict binds Glu (**Figure 3B**). We modelled a donor Glu into this pocket, positioning the :NH_2_ group near the acyl-adenylate for nucleophilic attack and formation of the Glu-Glu isopeptide bond. The γ-carboxyl was placed at a distance of ~3 Å from the SidJ R522 guanidinium, indicating a potential strong electrostatic interaction (**Figure 3C**). Mutation of R522 abolished SidJ-catalyzed glutamylation of SdeA (**Figure 3D**). Assuming the carboxyl group of a donor Asp side chain could enter the pocket and interact with R522, the distance between the :NH_2_ group of Asp and the acyl-adenylate would be too long for it to perform a nucleophilic attack. This mode of specificity is analogous to the Glu specificity observed for the tubulin glutamylase TTL6 (Mahalingan et al., 2020) (**Figure S6B**).

We also observed density in the kinase-like active site that corresponds to ATP and two Mg^2+^ ions (**Figure 3E**). The adenine is stabilized by stacking interaction with Y532 and the phosphates and Mg^2+^ ions are bound by kinase-like active site residues including K367 (PKA; K72), N534 (PKA; N171) and D542 (PKA; D184). Mutation of these residues reduced glutamylation activity (Black et al., 2019) (**Figure 3F**). Similar to canonical kinases, the ATP sits in a pocket between the two lobes of the kinase-like domain; however, the nucleotide is positioned in a unique manner. The β- and γ-phosphates of ATP are buried in a positively charged pocket, which is reminiscent of the pseudokinase SelO that binds ATP in a flipped orientation and transfers AMP from ATP to hydroxyl-containing amino acids (Sreelatha et al., 2018) (**Figure 3G**).

To determine the role of the kinase-like active site in SidJ, we performed adenylation reactions by monitoring ^32^P incorporation from [α-^32^P]ATP in the absence of Glu. The reactions were terminated with trichloroacetic acid (TCA) and adenylated SdeA^Core^ was detected as ^32^P-labelled protein in TCA-insoluble material. To account for any auto-modification of SidJ or non-specific modification of SdeA, we utilized a SdeA^Core^ E860Q mutant as a control and also added Glu to release the ^32^P-AMP signal from SdeA^Core^. Although the kinase-like active site residues are required for adenylation, His492 in the migrated nucleotide binding pocket is also required (**Figure 3H**), likely because it stabilizes the acyl-adenylate (**Figure 3A**). Notably, although the putative donor Glu-interacting Arg522 within the migrated nucleotide binding pocket is required for glutamylation (**Figure 3D**), the R522A mutant of SidJ still retains adenylation activity towards SdeA^Core^ (**Figure 3H**). Importantly, the addition of Glu to the R522A mutant reaction failed to decrease the ^32^P-signal in TCA precipitates. These results provide evidence that Arg522 in SidJ plays a major role in positioning the donor Glu. Taken together, the kinase-like active site of SidJ adenylates the SidE active site Glu, which is stabilized by the migrated nucleotide binding pocket to position the donor Glu for attack of the acyl-adenylate intermediate and the formation of an isopeptide bond.

### CaM Binding is Required for SidJ to Bind ATP and Adenylate SdeA

Several bacterial effectors and toxins are activated by eukaryote-specific proteins, such as CaM, to achieve spatial regulation (Guo et al., 2016; Guo et al., 2005; Leppla, 1984). CaM activates SidJ by binding to the IQ helix within the CTD (Black et al., 2019). However, the mechanism by which CaM activates SidJ is unknown. We monitored the thermal stability of SidJ and observed an ATP/Mg^2+^-dependent thermal shift that occurred only in the presence of CaM (**Figure 3I**). As expected, CaM was required for adenylation of SdeA^Core^ (**Figure 3J**). Thus, CaM binding renders SidJ competent to bind ATP and adenylate the SidE-ligases.

### The SidJ Paralog SdjA is an Active Glutamylating Enzyme

In *L. pneumophila*, the T4SS effector SdjA shares ~52% sequence identity to SidJ (**Figure 4A**); however, the function of SdjA is unknown. When overexpressed in yeast, SidJ, but not SdjA suppresses the growth inhibition phenotype induced by SdeA (Qiu et al., 2017a), suggesting different functions for the two effectors. Moreover, deletion of SidJ in *L. pneumophila* results in a growth inhibition phenotype that cannot be complemented by endogenous SdjA (Liu and Luo, 2007). Although the predicted kinase-like domains are 66% identical and the catalytic residues and CaM interacting IQ motif are well conserved, the NTDs are significantly different between SidJ and SdjA (**Figure S7A and B**). Because the majority of the interactions between SidJ and SdeA^Core^ lie within the SidJ NTD (**Figure 2B**), we asked whether SdjA could glutamylate the SidE-ligases. We incubated SdjA with CaM, ATP/Mg^2+^, [^14^C]-Glu and full-length SidE-family proteins and observed ^14^C incorporation into SdeB, SdeC and SidE, but not SdeA (**Figure 4B**). Consistent with our previous results (Black et al., 2019), SidJ glutamylated all 4 effectors. The SdjA reaction required CaM, ATP/Mg^2+^ and the residues in both the kinase-like active site and the migrated nucleotide binding pocket (**Figures 4C and D**). Similar to SidJ, SdjA glutamylated the active site Glu in SdeB, SdeC and SidE (**Figure 4E**). As expected, SdjA-mediated glutamylation completely abolished Ub ligase activity of SdeB, SdeC and SidE, in vitro (**Figure S7C**) and in cells (**Figure 4F**). Thus, SidJ and SdjA are CaM-dependent glutamylases that differentially inactivate the SidE-family of Ub ligases.

**Figure 4.**
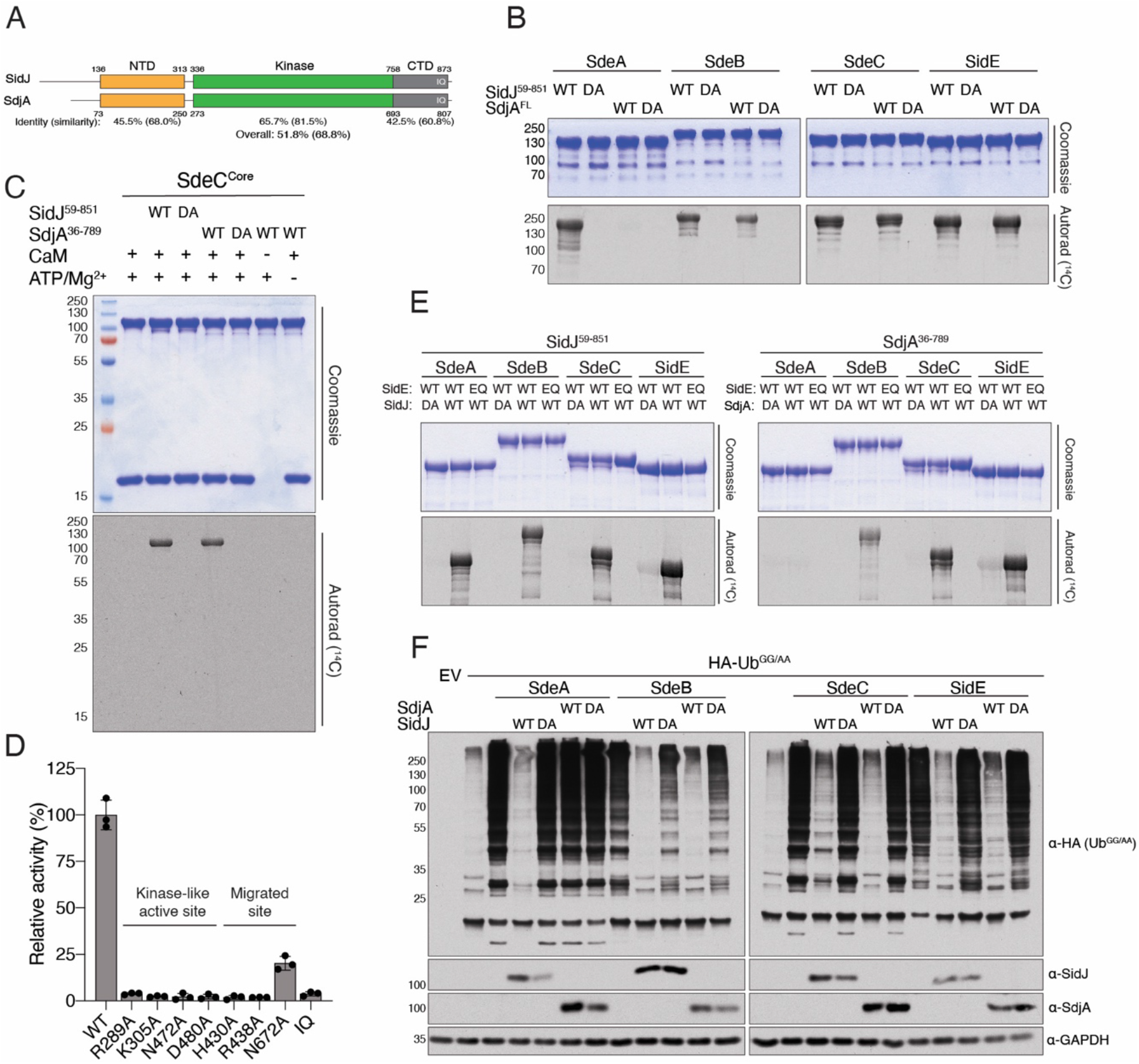
SidJ and SdjA differentially regulate the SidE-Ub ligases *in vitro*. **(A)** Cartoon representation of SidJ and SdjA depicting the NTD (orange), the kinase-like domain (green), and the CaM binding CTD (grey). The percent sequence identity and similarity are shown for each domain and for the overall protein. **(B)** Glutamylation activity of SidJ (SidJ^59-851^) and SdjA using full length SdeA, SdeB, SdeC and SidE as substrates. Reaction products were analyzed as in **Figure 2D**. DA denotes SidJ^D542A^ or SdjA^D480A^. Note that SdjA does not glutamylate SdeA. **(C)** Glutamylation activity of SidJ^59-851^ and SdjA^36-789^ using SdeC^Core^ as substrate. Reaction products were analyzed as in **Figure 2D**. DA denotes SidJ^D542A^ or SdjA^D480A^. **(D)** Glutamylation activity of SdjA^36-789^ and indicated mutants using SdeA^Core^ [^3^H]Glu as substrates. Reaction products were analyzed as in **Figure 2C**. The kinase-like active site residues and the residues in the migrated nucleotide binding pocket are highlighted. IQ mutant: SdjA^I776D; Q777D; R778E; R781E^. **(E)** Glutamylation activity of SidJ^59-851^ and SdjA^36-789^ using full length SdeA, SdeB, SdeC and SidE as substrates. Reaction products were analyzed as in **Figure 2D**. DA denotes SidJ^D542A^ or SdjA^D480A^. EQ denotes SdeA^E860Q^, SdeB^E857Q^, SdeC^E857Q^, or SidE^E855Q^. **(F)** Protein immunoblotting of total extracts from HEK293A cells expressing HA-Ub^GG/AA^, Myc-SdeA, Myc-SdeB, Myc-SdeC, or Myc-SidE and SidJ-V5, V5-SdjA or the indicated mutants. GAPDH is shown as a loading control. HA-Ub^GG/AA^ was used to specifically interrogate SidE-family activity because it cannot be used by the endogenous Ub machinery. DA denotes SidJ^D542A^ or SdjA^D480A^.

### The SidJ/SdjA NTDs Determine Specificity For the SidE Effectors

Because most of the interactions between SidJ/CaM with SdeA/C^Core^ come from the NTD, we hypothesized that the NTD could be the determining factor for the differential regulation of the SidE-family by SidJ and SdjA. We generated chimeric SidJ-SdjA proteins, in which the NTD from SidJ was replaced with the NTD from SdjA, and vice versa (**Figure 5A**). Replacing the HTH motif in SdjA with the corresponding HTH from SidJ resulted in an active chimeric protein that retained its ability to glutamylate SdeB, SdeC and SidE but also glutamylated SdeA (**Figure 5B, lane 9**). A SidJ chimera containing the SdjA NTD (SidJ^SdjA-NTD^) lost most of its activity towards SdeA (**Figure 5B, lane 13**). Thus, the variable NTD appears to be the major determinant of SidJ and SdjA specificity for the SidE effectors.

**Figure 5.**
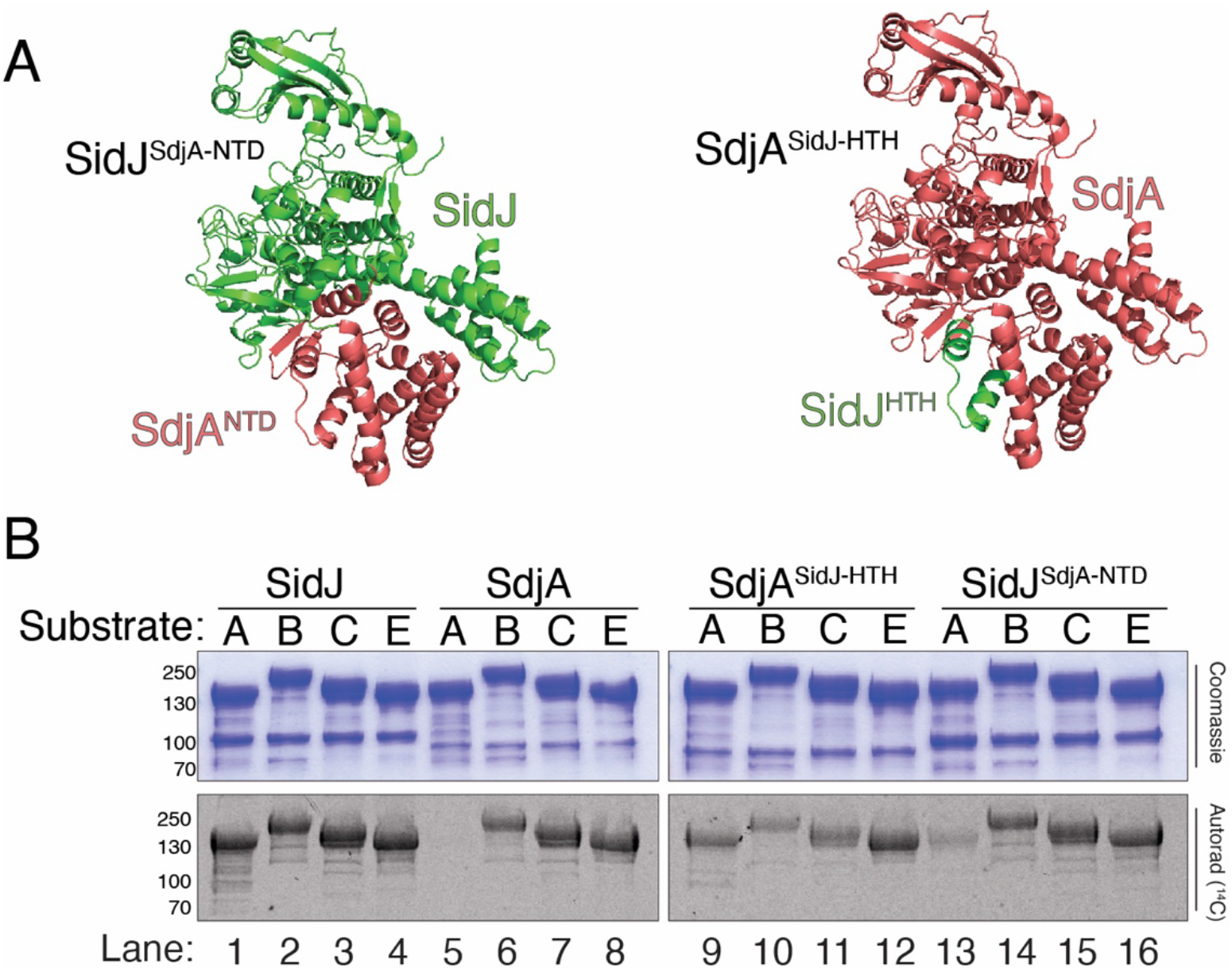
The variable NTD is the major determinant of SidJ and SdjA specificity for the SidE effectors. **(A)** Cartoon representation of SidJ^SdJA-NTD^ (left) and SdjA^SidJ-HTH^ (right) chimeras depicting the interchanged regions. SidJ is in green and SdjA is in salmon. **(B)** Glutamylation activity of SidJ^59-851^, SdjA, SdjA^SidJHTH^ and SidJ^SdJA-NTD^ using full length SdeA (A), SdeB (B), SdeC (C) and SidE (E) as substrates. Reaction products were analyzed as in **Figure 2D**. Compare lanes 1 and 13 (SidJ loss of activity), 5 and 9 (SdjA gain of activity).

### Differential Regulation of the SidE-effectors by SidJ and SdjA During *Legionella* Infection

SidJ is one of only a few T4SS effectors that when deleted from the *Legionella* genome causes a replication phenotype in infected cells, suggesting that SidJ and SdjA are not redundant (Jeong et al., 2015; Liu and Luo, 2007). We reasoned that in the background of a *sdeA* deletion, SidJ and SdjA would be functionally redundant (**Figure 6A**). We generated a *ΔsdjAΔsidJΔsdeA Legionella* strain and monitored its replication in the environmental host *Acanthamoeba castellanii* **(Figure S8)**. Remarkably, both SidJ and SdjA could complement the growth defect in the *ΔsdjAΔsidJΔsdeA* strain (**Figure 6B**). In contrast, SidJ, but not SdjA could rescue the growth defect in a *ΔsdjAΔsidJ* strain (**Figure 6C**). Collectively, our results suggest that the SidE-Ub ligases are differentially regulated by SidJ and SdjA during *Legionella* infection.

**Figure 6.**
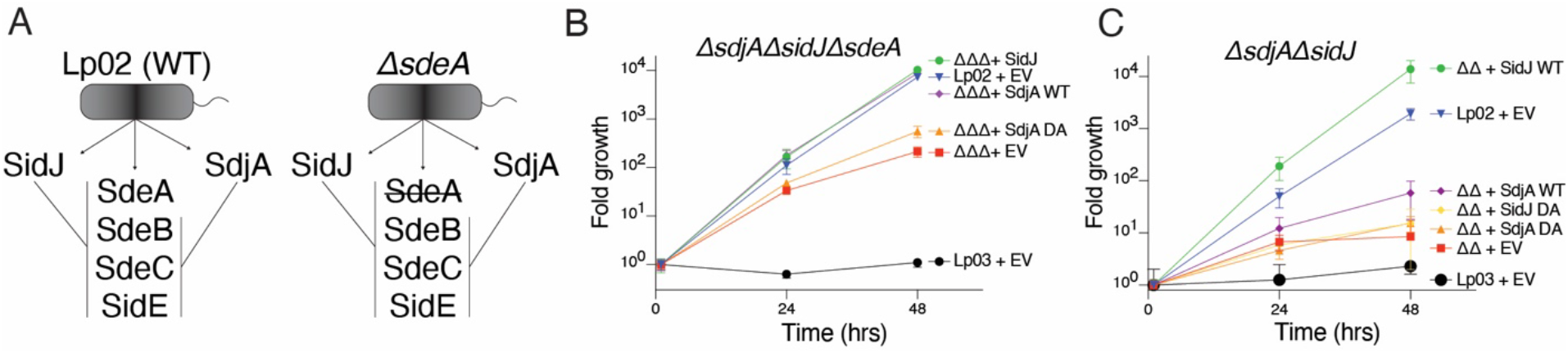
SidJ and SdjA differentially regulate the SidE-Ub ligases during *Legionella* infection. **(A)** Schematic representation of SidJ/SdjA genetic interaction during *Legionella* infection. In a wild-type *L. pneumophila* (Lp02) strain, SdjA is unable to complement SidJ because it is inactive against SdeA (left). In a *ΔsdeA* background, SdjA is functionally redundant with SidJ (right). **(B, C)** Replication of *L. pneumophila* strains in *A. castellanii.* Infected amoeba cells were lysed at the indicated timepoints and bacterial replication was quantified by plating serial dilutions of lysates. Results are representative of two independent experiments; error bars denote STD from 1 experiment performed in triplicate. *ΔsdjAΔsidJΔsdeA* (ΔΔΔ), *ΔsidJΔsdeA* (ΔΔ), EV; empty vector, DA denotes SidJ^D542A^ or SdjA^D480A^, Lp03; *ΔdotA* (T4SS deficient).

## Discussion

We propose a model for how SidJ glutamylates the SidE-family of Ub ligases (**Figure 7**). During infection, SidJ is translocated into the host cell where it binds CaM (1), which allows the kinase-like active site to bind ATP/Mg^2+^ (2). Notably, because CaM is a eukaryote-specific protein, this mechanism allows for spatial regulation of the SidE-Ub ligases within the host cell. SidJ then adenylates the active site Glu within the ARTT loop in the SidE effectors (3). Adenylated SidE binds the migrated nucleotide binding pocket (4), which positions free Glu near the acyl-adenylate intermediate for nucleophilic attack and subsequent formation of the Glu-Glu isopeptide bond (5).

**Figure 7.**
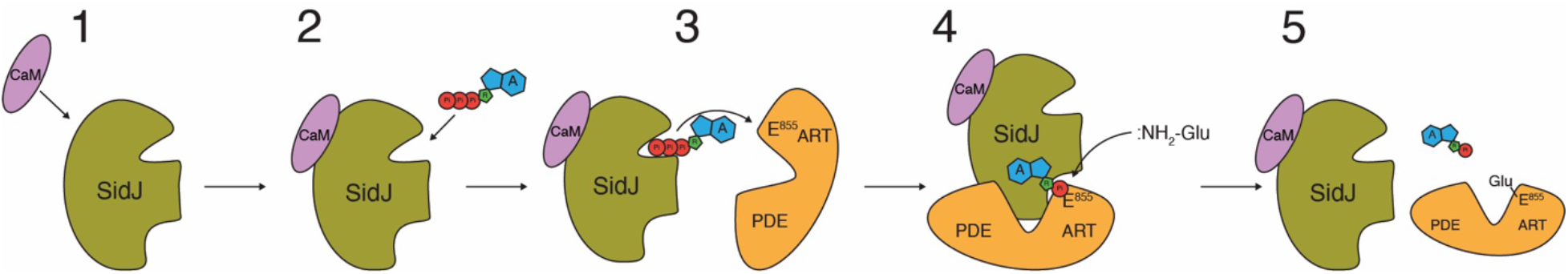
Model of SidJ-catalyzed glutamylation of the SidE effectors. **(1)** SidJ is translocated into host cells and binds CaM. **(2)** CaM binding renders the kinase-like active site competent to bind ATP/Mg^2+^ (**Figure 3I**) and **(3)** adenylate the active site Glu in the SidE effectors (**Figures 3E-H**). **(4)** Adenylated SidE binds the migrated nucleotide binding pocket of SidJ, forming a stable reaction intermediate (**Figures 1D and 2A**). Free Glu binds to a positively charged cleft in the migrated nucleotide binding pocket of SidJ (**Figures 3B-D**), which positions the NH_2_ group for nucleophilic attack of the acyl-adenylate, **(5)** releasing AMP and forming an isopeptide bond (**Figure 1C**).

Our structural and biochemical data suggest that the kinase-like active site performs the adenylation reaction and the migrated nucleotide binding pocket executes the glutamylation reaction. Although His492 in the migrated nucleotide binding pocket is required for adenylation and glutamylation, the R522A SidJ mutant retains adenylation activity, fails to bind the donor Glu and is inactive in glutamylation reactions. Because Arg522 does not contact the acyl-adenylate, His492 appears to be required for adenylation because it stabilizes the acyl-adenylate intermediate (**Figure 3A**).

SidJ homologs are found in taxonomically diverse groups of organisms including archaea and viruses, suggesting that SidJ is being spread by horizontal gene transfer (**Figure S9A**). None of these organisms except for *Legionella* species have SidE homologs or the SidJ CaM-binding IQ motif and they show virtually no conservation within the SidE-interacting NTD (**Figure S9B**). Thus, although the kinase-like active site residues are conserved among SidJ homologs, their substrates and activation mechanisms will likely differ. Interestingly, SidJ homologs from crocodilepox viruses are located next to a cluster of proteins that differ from SidE but are involved in the modulation of host ubiquitination pathways (Afonso et al., 2006).

In mammals, tubulin glutamylation, tyrosination and glycylation are performed by Tubulin-tyrosine ligase-like enzymes (TTLL), members of the ATP-grasp superfamily (Song and Brady, 2015; Yu et al., 2015). TTLLs catalyze amino acid ligation via the formation of a high-energy acyl-phosphate intermediate, which is subsequently attacked by the amine group of the donor amino acid (Szyk et al., 2011). Interestingly, both the protein kinase and ATP-grasp fold enzymes share a common topology (Grishin, 1999). In contrast to most protein kinases, SidJ adenylates a carboxyl containing amino acid as an intermediate step in peptide bond formation. We propose that the protein kinase fold can phosphorylate or adenylate carboxylic groups to mediate non-ribosomal amino-acid ligation reactions. Notably, a recent study of “the hidden phosphoproteome” revealed that a vast number of Glu and Asp residues within HeLa cells are phosphorylated (Hardman et al., 2019).

In summary, our work underscores the catalytic versatility of the kinase fold and reveals a previously unappreciated level of regulation of the SidE Ub ligases.

## Acknowledgments

We thank Diana Tomchick and members of the Tagliabracci laboratory for discussions. Results shown are derived from work performed at the Pacific Northwest Center for Cryo-EM. This work was funded by NIH Grants DP2GM137419 (V.S.T.), F30HL143859 (M.H.B.), a W. M. Keck Foundation grant (V.S.T and K.P.), Welch Foundation Grant I-1911 (V.S.T.), Polish National Agency for Scientific Exchange scholarship PPN/BEK/2018/1/00431 (K.P.). We thank the Structural Biology Laboratory and the Cryo Electron Microscopy Facility at UT Southwestern Medical Center which are partially supported by grant RP170644 from the Cancer Prevention & Research Institute of Texas (CPRIT) for cryo-EM studies. A portion of this research was supported by NIH grant U24GM129547 and performed at the PNCC at OHSU and accessed through EMSL (grid.436923.9), a DOE Office of Science User Facility sponsored by the Office of Biological and Environmental Research. V.S.T. is a Michael L. Rosenberg Scholar in Medical Research, a CPRIT Scholar (RR150033) and a Searle Scholar.

## Author Contributions

A.O., M.H.B., Z.C., Y.L. and V.S.T. designed the experiments. A.O., M.H.B., Z.C., Y.L. and V.S.T. conducted the experiments. A.O., Z.C., Y.L. performed the cryo-EM. K.P. performed the bioinformatics. A.O., K.P. and V.S.T. wrote the manuscript with input from all authors.

## References

Adams, P.D., Afonine, P.V., Bunkoczi, G., Chen, V.B., Davis, I.W., Echols, N., Headd, J.J., Hung, L.W., Kapral, G.J., Grosse-Kunstleve, R.W., et al. (2010). PHENIX: a comprehensive Python-based system for macromolecular structure solution. Acta Crystallogr D Biol Crystallogr 66, 213-221.

Afonso, C.L., Tulman, E.R., Delhon, G., Lu, Z., Viljoen, G.J., Wallace, D.B., Kutish, G.F., and Rock, D.L. (2006). Genome of crocodilepox virus. J Virol 80, 4978-4991.

Akturk, A., Wasilko, D.J., Wu, X., Liu, Y., Zhang, Y., Qiu, J., Luo, Z.Q., Reiter, K.H., Brzovic, P.S., Klevit, R.E., et al. (2018). Mechanism of phosphoribosyl-ubiquitination mediated by a single Legionella effector. Nature 557, 729-733.

Bardill, J.P., Miller, J.L., and Vogel, J.P. (2005). IcmS-dependent translocation of SdeA into macrophages by the Legionella pneumophila type IV secretion system. Mol Microbiol 56, 90-103.

Bhogaraju, S., Bonn, F., Mukherjee, R., Adams, M., Pfleiderer, M.M., Galej, W.P., Matkovic, V., Lopez-Mosqueda, J., Kalayil, S., Shin, D., et al. (2019). Inhibition of bacterial ubiquitin ligases by SidJ-calmodulin catalysed glutamylation. Nature 572, 382-386.

Bhogaraju, S., Kalayil, S., Liu, Y., Bonn, F., Colby, T., Matic, I., and Dikic, I. (2016). Phosphoribosylation of Ubiquitin Promotes Serine Ubiquitination and Impairs Conventional Ubiquitination. Cell 167, 1636-1649 e1613.

Black, M.H., Osinski, A., Gradowski, M., Servage, K.A., Pawlowski, K., Tomchick, D.R., and Tagliabracci, V.S. (2019). Bacterial pseudokinase catalyzes protein polyglutamylation to inhibit the SidE-family ubiquitin ligases. Science 364, 787-792.

Chen, V.B., Arendall, W.B., 3rd, Headd, J.J., Keedy, D.A., Immormino, R.M., Kapral, G.J., Murray, L.W., Richardson, J.S., and Richardson, D.C. (2010). MolProbity: all-atom structure validation for macromolecular crystallography. Acta Crystallogr D Biol Crystallogr 66, 12-21.

Cornejo, E., Schlaermann, P., and Mukherjee, S. (2017). How to rewire the host cell: A home improvement guide for intracellular bacteria. J Cell Biol 216, 3931-3948.

Crooks, G.E., Hon, G., Chandonia, J.M., and Brenner, S.E. (2004). WebLogo: a sequence logo generator. Genome Res 14, 1188-1190.

Dereeper, A., Guignon, V., Blanc, G., Audic, S., Buffet, S., Chevenet, F., Dufayard, J.F., Guindon, S., Lefort, V., Lescot, M., et al. (2008). Phylogeny.fr: robust phylogenetic analysis for the non-specialist. Nucleic Acids Res 36, W465-469.

Dong, Y., Mu, Y., Xie, Y., Zhang, Y., Han, Y., Zhou, Y., Wang, W., Liu, Z., Wu, M., Wang, H., et al. (2018). Structural basis of ubiquitin modification by the Legionella effector SdeA. Nature 557, 674-678.

Emsley, P., Lohkamp, B., Scott, W.G., and Cowtan, K. (2010). Features and development of Coot. Acta Crystallogr D Biol Crystallogr 66, 486-501.

Gan, N., Zhen, X., Liu, Y., Xu, X., He, C., Qiu, J., Liu, Y., Fujimoto, G.M., Nakayasu, E.S., Zhou, B., et al. (2019). Regulation of phosphoribosyl ubiquitination by a calmodulin-dependent glutamylase. Nature 572, 387-391.

Garnham, C.P., Vemu, A., Wilson-Kubalek, E.M., Yu, I., Szyk, A., Lander, G.C., Milligan, R.A., and Roll-Mecak, A. (2015). Multivalent Microtubule Recognition by Tubulin Tyrosine Ligase-like Family Glutamylases. Cell 161, 1112-1123.

Grishin, N.V. (1999). Phosphatidylinositol phosphate kinase: a link between protein kinase and glutathione synthase folds. J Mol Biol 291, 239-247.

Guo, M., Kim, P., Li, G., Elowsky, C.G., and Alfano, J.R. (2016). A Bacterial Effector Co-opts Calmodulin to Target the Plant Microtubule Network. Cell Host Microbe 19, 67-78.

Guo, Q., Shen, Y., Lee, Y.S., Gibbs, C.S., Mrksich, M., and Tang, W.J. (2005). Structural basis for the interaction of Bordetella pertussis adenylyl cyclase toxin with calmodulin. EMBO J 24, 3190-3201.

Hardman, G., Perkins, S., Brownridge, P.J., Clarke, C.J., Byrne, D.P., Campbell, A.E., Kalyuzhnyy, A., Myall, A., Eyers, P.A., Jones, A.R., et al. (2019). Strong anion exchange-mediated phosphoproteomics reveals extensive human non-canonical phosphorylation. EMBO J 38, e100847.

Havey, J.C., and Roy, C.R. (2015). Toxicity and SidJ-Mediated Suppression of Toxicity Require Distinct Regions in the SidE Family of Legionella pneumophila Effectors. Infect Immun 83, 3506-3514.

Huang, Y., Niu, B., Gao, Y., Fu, L., and Li, W. (2010). CD-HIT Suite: a web server for clustering and comparing biological sequences. Bioinformatics 26, 680-682.

Isberg, R.R., O’Connor, T.J., and Heidtman, M. (2009). The Legionella pneumophila replication vacuole: making a cosy niche inside host cells. Nat Rev Microbiol 7, 13-24.

Jeong, K.C., Sexton, J.A., and Vogel, J.P. (2015). Spatiotemporal regulation of a Legionella pneumophila T4SS substrate by the metaeffector SidJ. PLoS Pathog 11, e1004695.

Kalayil, S., Bhogaraju, S., Bonn, F., Shin, D., Liu, Y., Gan, N., Basquin, J., Grumati, P., Luo, Z.Q., and Dikic, I. (2018). Insights into catalysis and function of phosphoribosyl-linked serine ubiquitination. Nature 557, 734-738.

Katoh, K., Rozewicki, J., and Yamada, K.D. (2019). MAFFT online service: multiple sequence alignment, interactive sequence choice and visualization. Brief Bioinform 20, 1160-1166.

Kent, U.M. (1999). Purification of antibodies using ammonium sulfate fractionation or gel filtration. Methods Mol Biol 115, 11-18.

Kotewicz, K.M., Ramabhadran, V., Sjoblom, N., Vogel, J.P., Haenssler, E., Zhang, M., Behringer, J., Scheck, R.A., and Isberg, R.R. (2017). A Single Legionella Effector Catalyzes a Multistep Ubiquitination Pathway to Rearrange Tubular Endoplasmic Reticulum for Replication. Cell Host Microbe 21, 169-181.

Kwon, A., Scott, S., Taujale, R., Yeung, W., Kochut, K.J., Eyers, P.A., and Kannan, N. (2019). Tracing the origin and evolution of pseudokinases across the tree of life. Sci Signal 12.

Leppla, S.H. (1984). Bacillus anthracis calmodulin-dependent adenylate cyclase: chemical and enzymatic properties and interactions with eucaryotic cells. Adv Cyclic Nucleotide Protein Phosphorylation Res 17, 189-198.

Letunic, I., and Bork, P. (2016). Interactive tree of life (iTOL) v3: an online tool for the display and annotation of phylogenetic and other trees. Nucleic Acids Res 44, W242-245.

Li, Y., Cash, J.N., Tesmer, J.J.G., and Cianfrocco, M.A. (2020). High-Throughput Cryo-EM Enabled by User-Free Preprocessing Routines. Structure 28, 858-869 e853.

Liu, Y., and Luo, Z.Q. (2007). The Legionella pneumophila effector SidJ is required for efficient recruitment of endoplasmic reticulum proteins to the bacterial phagosome. Infect Immun 75, 592-603.

Mahalingan, K.K., Keith Keenan, E., Strickland, M., Li, Y., Liu, Y., Ball, H.L., Tanner, M.E., Tjandra, N., and Roll-Mecak, A. (2020). Structural basis for polyglutamate chain initiation and elongation by TTLL family enzymes. Nat Struct Mol Biol 27, 802-813.

Mastronarde, D.N. (2005). Automated electron microscope tomography using robust prediction of specimen movements. J Struct Biol 152, 36-51.

Mukherjee, S., Liu, X., Arasaki, K., McDonough, J., Galan, J.E., and Roy, C.R. (2011). Modulation of Rab GTPase function by a protein phosphocholine transferase. Nature 477, 103-106.

Neunuebel, M.R., Chen, Y., Gaspar, A.H., Backlund, P.S., Jr., Yergey, A., and Machner, M.P. (2011). De-AMPylation of the small GTPase Rab1 by the pathogen *Legionella pneumophila*. Science 333, 453–456.

Pettersen, E.F., Goddard, T.D., Huang, C.C., Couch, G.S., Greenblatt, D.M., Meng, E.C., and Ferrin, T.E. (2004). UCSF Chimera--a visualization system for exploratory research and analysis. J Comput Chem 25, 1605-1612.

Qiu, J., Sheedlo, M.J., Yu, K., Tan, Y., Nakayasu, E.S., Das, C., Liu, X., and Luo, Z.Q. (2016). Ubiquitination independent of E1 and E2 enzymes by bacterial effectors. Nature 533, 120-124.

Qiu, J., Yu, K., Fei, X., Liu, Y., Nakayasu, E.S., Piehowski, P.D., Shaw, J.B., Puvar, K., Das, C., Liu, X., et al. (2017a). A unique deubiquitinase that deconjugates phosphoribosyl-linked protein ubiquitination. Cell Research 27, 865-881.

Qiu, J., Yu, K., Fei, X., Liu, Y., Nakayasu, E.S., Piehowski, P.D., Shaw, J.B., Puvar, K., Das, C., Liu, X., et al. (2017b). A unique deubiquitinase that deconjugates phosphoribosyl-linked protein ubiquitination. Cell Res 27, 865-881.

Ribeiro, A.J.M., Das, S., Dawson, N., Zaru, R., Orchard, S., Thornton, J.M., Orengo, C., Zeqiraj, E., Murphy, J.M., and Eyers, P.A. (2019). Emerging concepts in pseudoenzyme classification, evolution, and signaling. Sci Signal 12.

Robert, X., and Gouet, P. (2014). Deciphering key features in protein structures with the new ENDscript server. Nucleic Acids Res 42, W320-324.

Sanchez-Garcia, R., Gomez-Blanco, J., Cuervo, A., Carazo, J., Sorzano, C., and Vargas, J. (2020). DeepEMhancer: a deep learning solution for cryo-EM volume post-processing. bioRxiv, 2020.2006.2012.148296.

Scheres, S.H. (2012). RELION: implementation of a Bayesian approach to cryo-EM structure determination. J Struct Biol 180, 519-530.

Shin, D., Mukherjee, R., Liu, Y., Gonzalez, A., Bonn, F., Liu, Y., Rogov, V.V., Heinz, M., Stolz, A., Hummer, G., et al. (2020). Regulation of Phosphoribosyl-Linked Serine Ubiquitination by Deubiquitinases DupA and DupB. Mol Cell 77, 164-179 e166.

Song, Y., and Brady, S.T. (2015). Post-translational modifications of tubulin: pathways to functional diversity of microtubules. Trends Cell Biol 25, 125-136.

Sreelatha, A., Yee, S.S., Lopez, V.A., Park, B.C., Kinch, L.N., Pilch, S., Servage, K.A., Zhang, J., Jiou, J., Karasiewicz-Urbanska, M., et al. (2018). Protein AMPylation by an Evolutionarily Conserved Pseudokinase. Cell 175, 809-821 e819.

Sulpizio, A., Minelli, M.E., Wan, M., Burrowes, P.D., Wu, X., Sanford, E.J., Shin, J.H., Williams, B.C., Goldberg, M.L., Smolka, M.B., et al. (2019). Protein polyglutamylation catalyzed by the bacterial calmodulin-dependent pseudokinase SidJ. eLife 8.

Szyk, A., Deaconescu, A.M., Piszczek, G., and Roll-Mecak, A. (2011). Tubulin tyrosine ligase structure reveals adaptation of an ancient fold to bind and modify tubulin. Nat Struct Mol Biol 18, 1250-1258.

Wagner, T., Merino, F., Stabrin, M., Moriya, T., Antoni, C., Apelbaum, A., Hagel, P., Sitsel, O., Raisch, T., Prumbaum, D., et al. (2019). SPHIRE-crYOLO is a fast and accurate fully automated particle picker for cryo-EM. Commun Biol 2, 218.

Wan, M., Sulpizio, A.G., Akturk, A., Beck, W.H.J., Lanz, M., Faca, V.M., Smolka, M.B., Vogel, J.P., and Mao, Y. (2019). Deubiquitination of phosphoribosyl-ubiquitin conjugates by phosphodiesterase-domain-containing Legionella effectors. Proc Natl Acad Sci U S A 116, 23518-23526.

Wang, Y., Shi, M., Feng, H., Zhu, Y., Liu, S., Gao, A., and Gao, P. (2018). Structural Insights into Non-canonical Ubiquitination Catalyzed by SidE. Cell 173, 1231-1243 e1216.

Winn, M.D., Ballard, C.C., Cowtan, K.D., Dodson, E.J., Emsley, P., Evans, P.R., Keegan, R.M., Krissinel, E.B., Leslie, A.G., McCoy, A., et al. (2011). Overview of the CCP4 suite and current developments. Acta Crystallogr D Biol Crystallogr 67, 235-242.

Yu, I., Garnham, C.P., and Roll-Mecak, A. (2015). Writing and Reading the Tubulin Code. J Biol Chem 290, 17163-17172.

Zhang, K. (2016). Gctf: Real-time CTF determination and correction. J Struct Biol 193, 1-12.

Zheng, S.Q., Palovcak, E., Armache, J.P., Verba, K.A., Cheng, Y., and Agard, D.A. (2017). MotionCor2: anisotropic correction of beam-induced motion for improved cryo-electron microscopy. Nat Methods 14, 331-332.

