## Supplemental Materials for "Structural and mechanistic basis for protein glutamylation by the kinase fold"

**Supplemental Figure Titles and Legends**

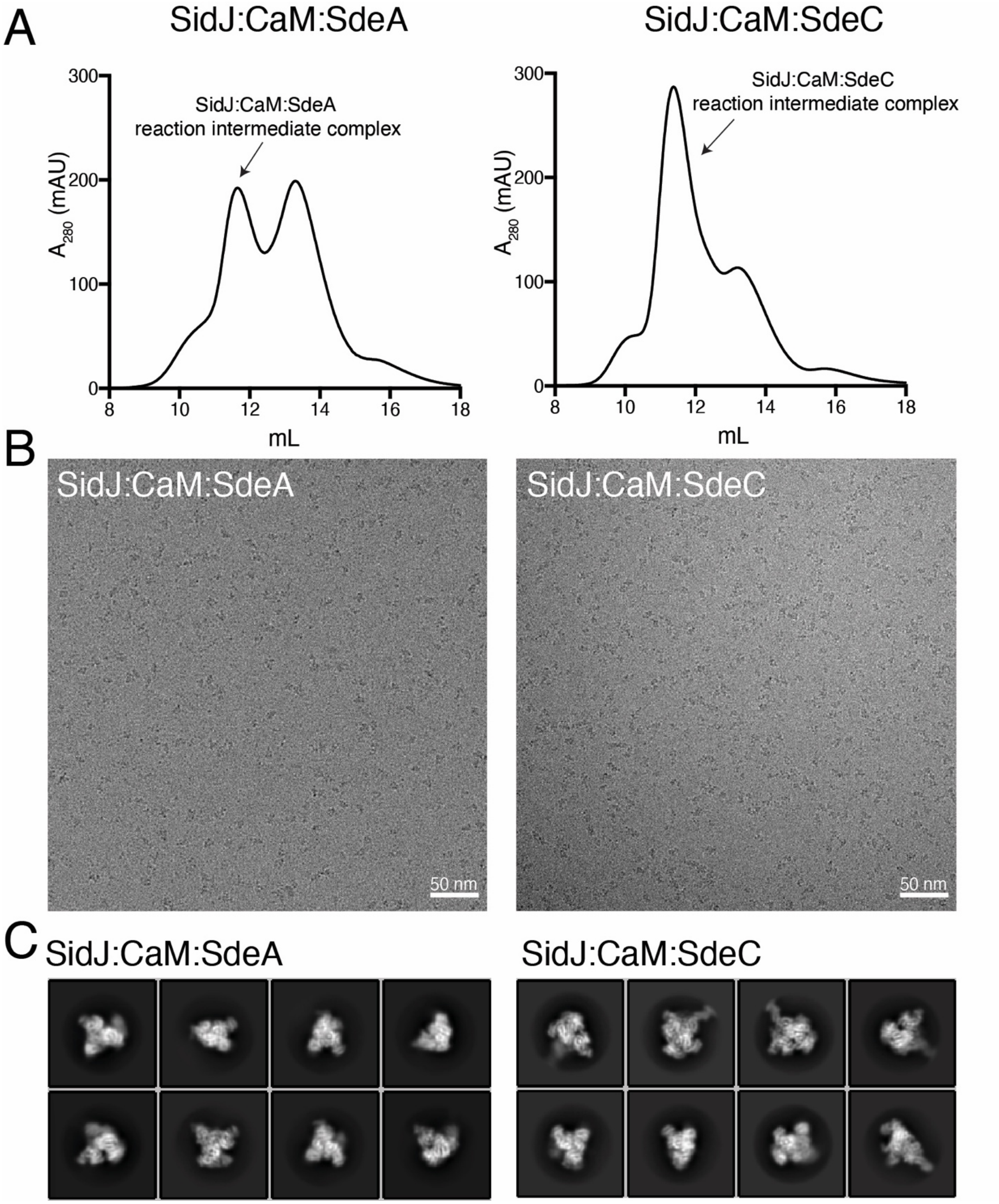

**Figure S1. Cryo-EM analysis of the SidJ:CaM:SdeA and SidJ:CaM:SdeC reaction intermediate complexes**.

**(A)** SEC traces of SidJ:CaM:SdeA (left) and SidJ:CaM:SdeC (right) reaction intermediate complexes.

**(B)** Representative micrographs depicting the SidJ:CaM:SdeA (left) and SidJ:CaM:SdeC (right) reaction intermediate complexes.

**(C)** Representative 2D classes generated by Relion 2D-classification for the SidJ:CaM:SdeA (left) and SidJ:CaM:SdeC (right) reaction intermediate complexes.

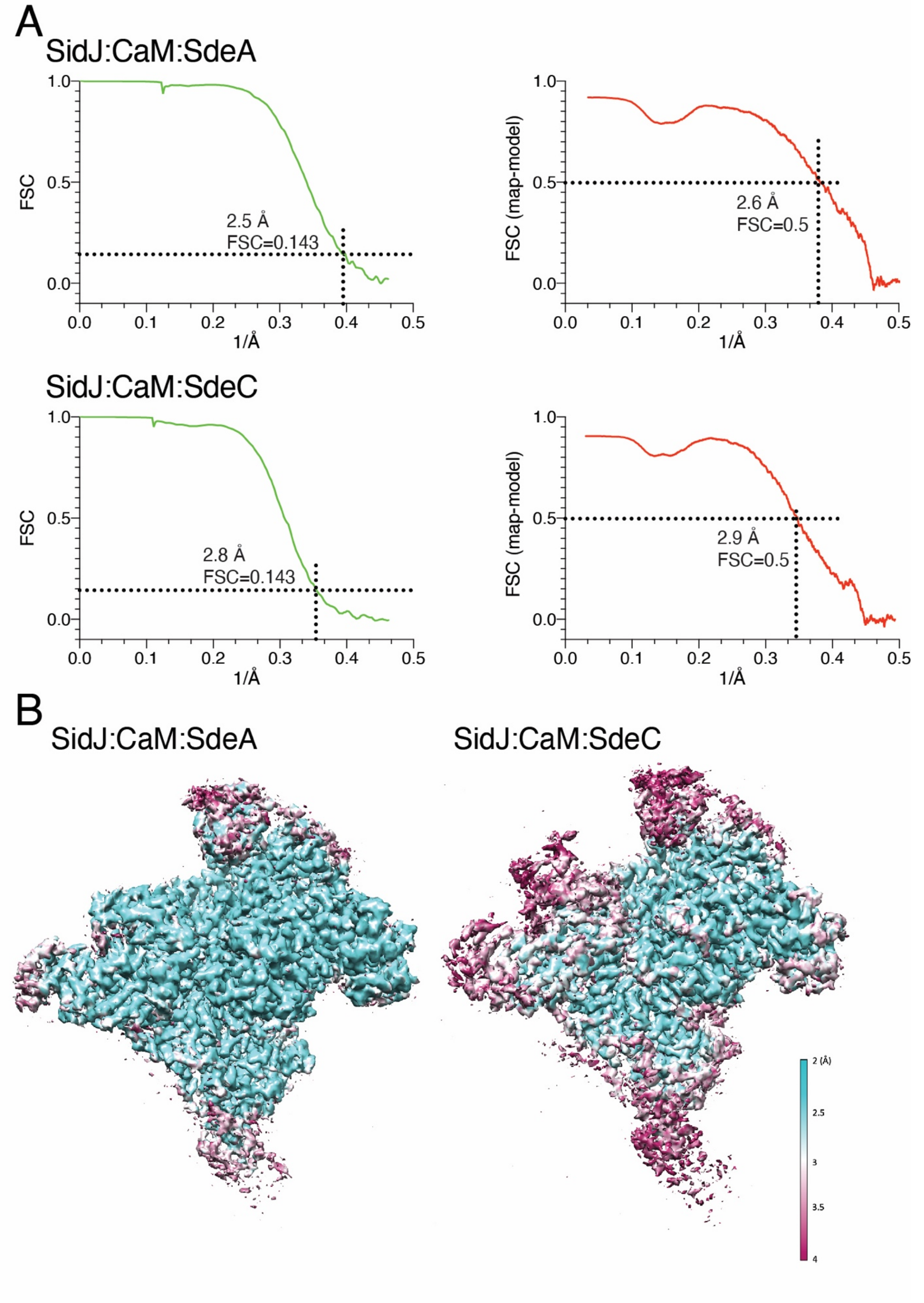
A
SidJ:CaM:SdeA
FSC
1/Å
2.5 Å
FSC=0.143
FSC (map-model)
2.6 Å
FSC=0.5
SidJ:CaM:SdeC
2.8 Å
FSC=0.143
2.9 Å
FSC=0.5
B
SidJ:CaM:SdeA
SidJ:CaM:SdeC
2 (Å)
2.5
3
3.5
4

**Figure S2. Cryo-EM analysis of the SidJ:CaM:SdeA and SidJ:CaM:SdeC reaction intermediate complexes**.

**(A)** Gold-standard FSC curves (green trace) of SidJ:CaM:SdeA (upper) and SidJ:CaM:SdeC (lower) and corresponding map-model FSC curves (red curves).

**(B)** Local resolution of the SidJ:CaM:SdeA (left) and SidJ:CaM:SdeC (right) reaction intermediate complexes calculated by RELION.

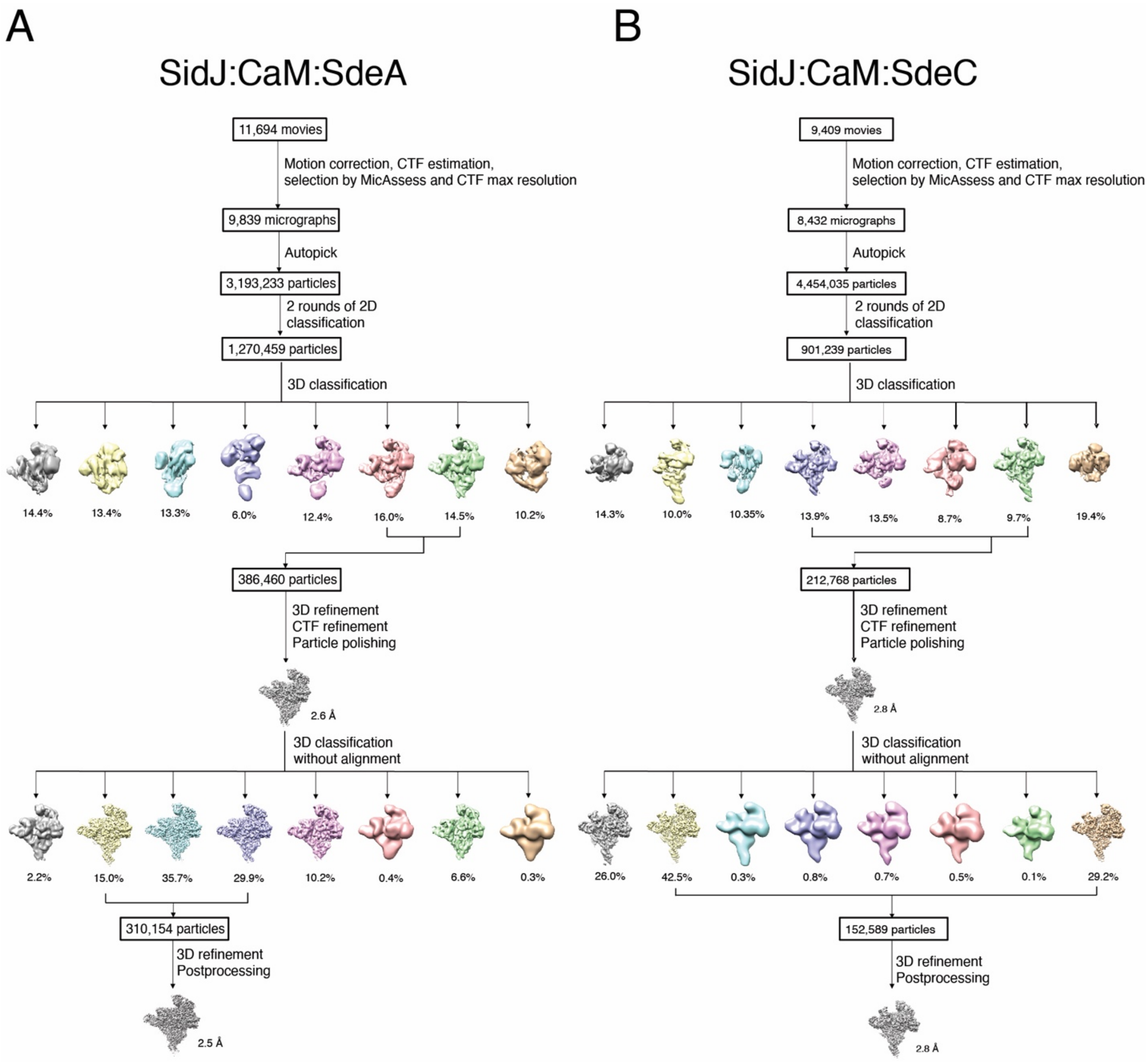


**Figure S3**. **Flow chart of image processing.**

**(A,B)** Flow charts depicting the image processing of the SidJ:CaM:SdeA (**A**) and SidJ:CaM:SdeC (**B**) reaction intermediate complexes.

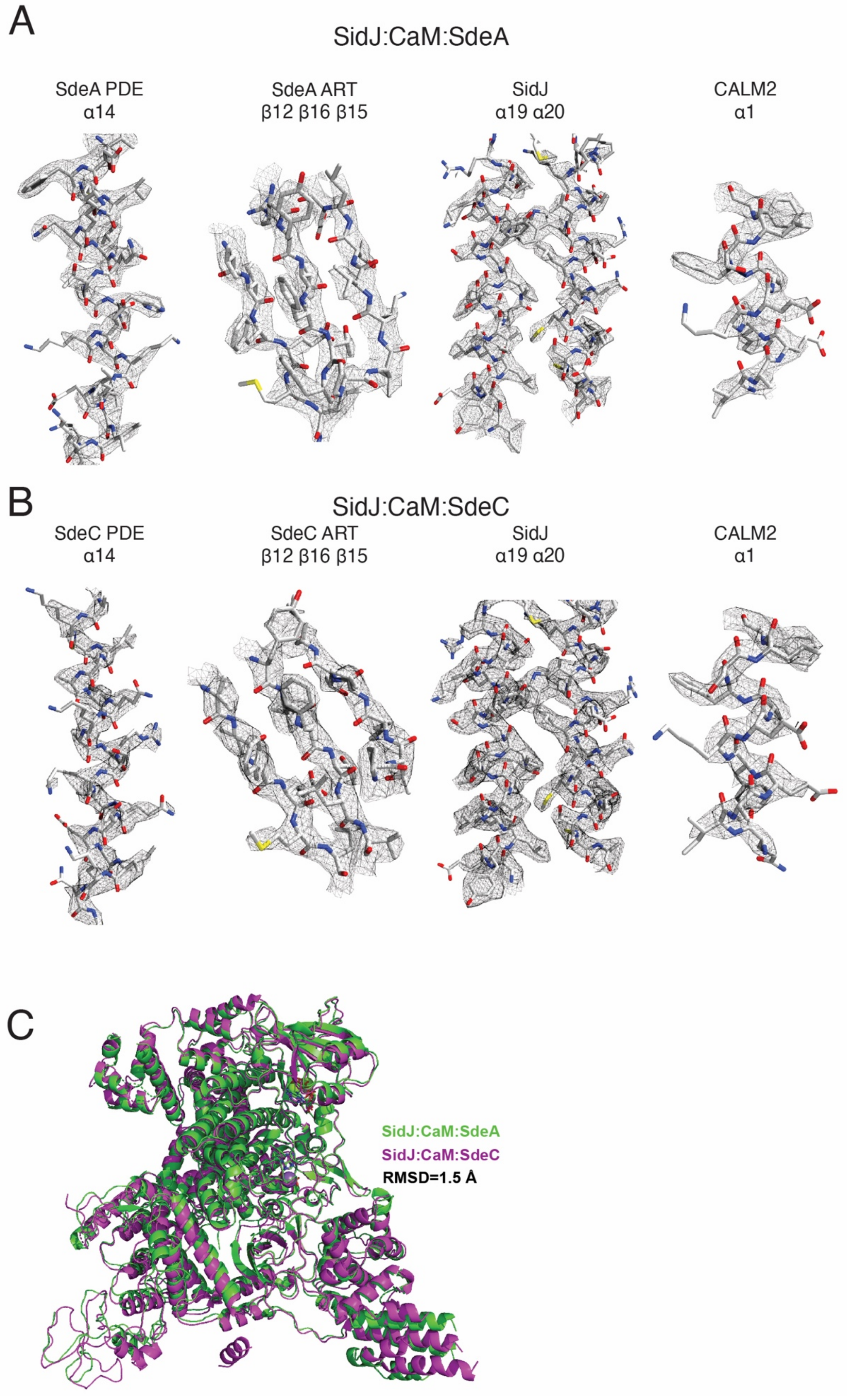
A
SidJ:CaM:SdeA
SdeA PDE
α14
SdeA ART
β12 β16 β15
SidJ
α19 α20
CALM2
α1
B
SidJ:CaM:SdeC
SdeC PDE
α14
SdeC ART
β12 β16 β15
SidJ
α19 α20
CALM2
α1
C
SidJ:CaM:SdeA
SidJ:CaM:SdeC
RMSD=1.5 Å

**Figure S4. Representative density maps of SidJ, SdeA, SdeC and CaM within indicated complexes.**

**(A)** Density maps for the SidJ:CaM:SdeA structure. Models fit into the density are shown as white sticks.

**(B)** Density maps for the SidJ:CaM:SdeC structure.

**(C)** Structural comparison of the SidJ:CaM:SdeA (green) and SidJ:CaM:SdeC (purple) reaction intermediate complexes. Ribbon diagrams showing the aligned structures of the two complexes, with a RMSD of 1.5 Å.

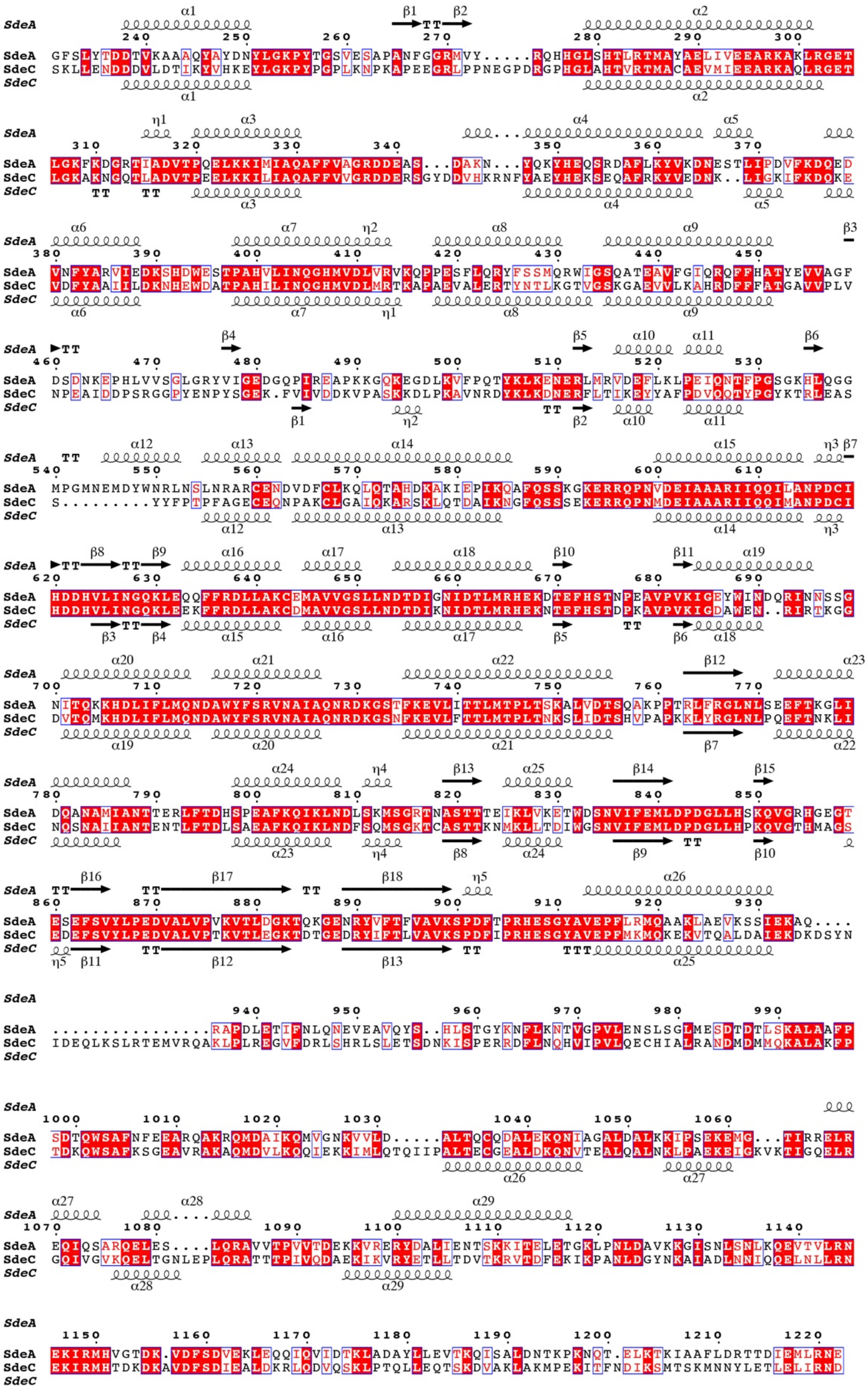

**Figure S5**. **Pairwise sequence alignment of SdeA$^{\text{Core}}$ and SdeC$^{\text{Core}}$.** Secondary structure was annotated from the cryo-EM models. Alignment figure was generated by ESPript 3 (Robert and Gouet, 2014).

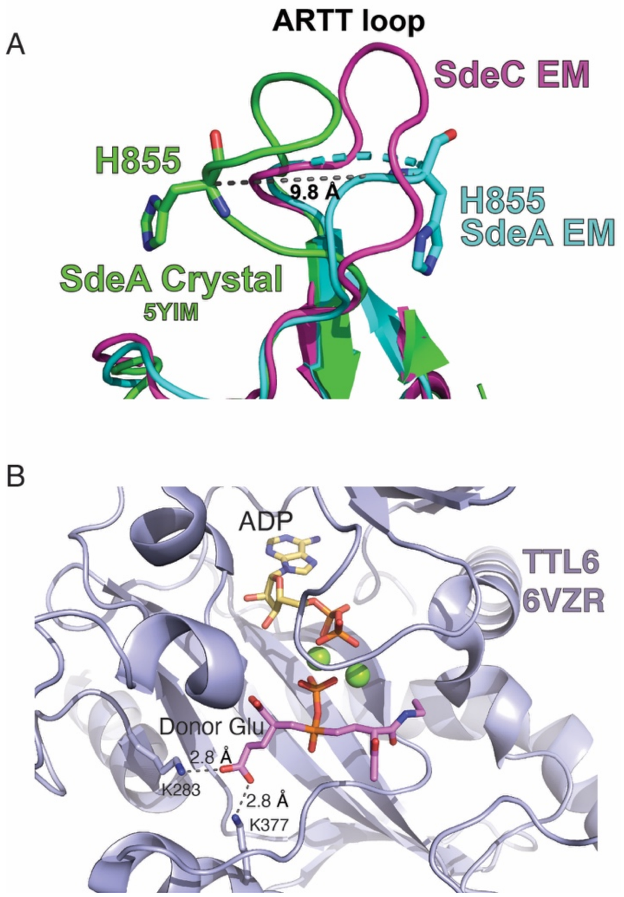


**Figure S6**. **Unique active site architectures mediate acyl-adenylate formation and glutamylation.**

**(A)** Conformational changes in the SdeA ARTT loop upon SidJ binding and acyl-adenylate formation. The ARTT loop of SdeA in complex with SidJ changes its conformation, as compared to the crystal structure of apo-SdeA (PDBID: 5YIM). The 9.8 Å loop movement is measured as a distance between Cα of H855 in SdeA crystal structure (green), and cryo-EM complex with SdeA (cyan).

**(B)** The structural basis for Tubulin Tyrosinase-Like 6 (TTL6) glutamylase specificity for Glu may be similar to SidJ. Details of the active site of TTL6 (pale blue) depicting a phosphorylated initiation analog in stick (PDBID: 6VZR). The initiation analogue (magenta) has a γ-carboxyl of its donor Glu bound by K283 and K377, in optimal hydrogen bonding length.

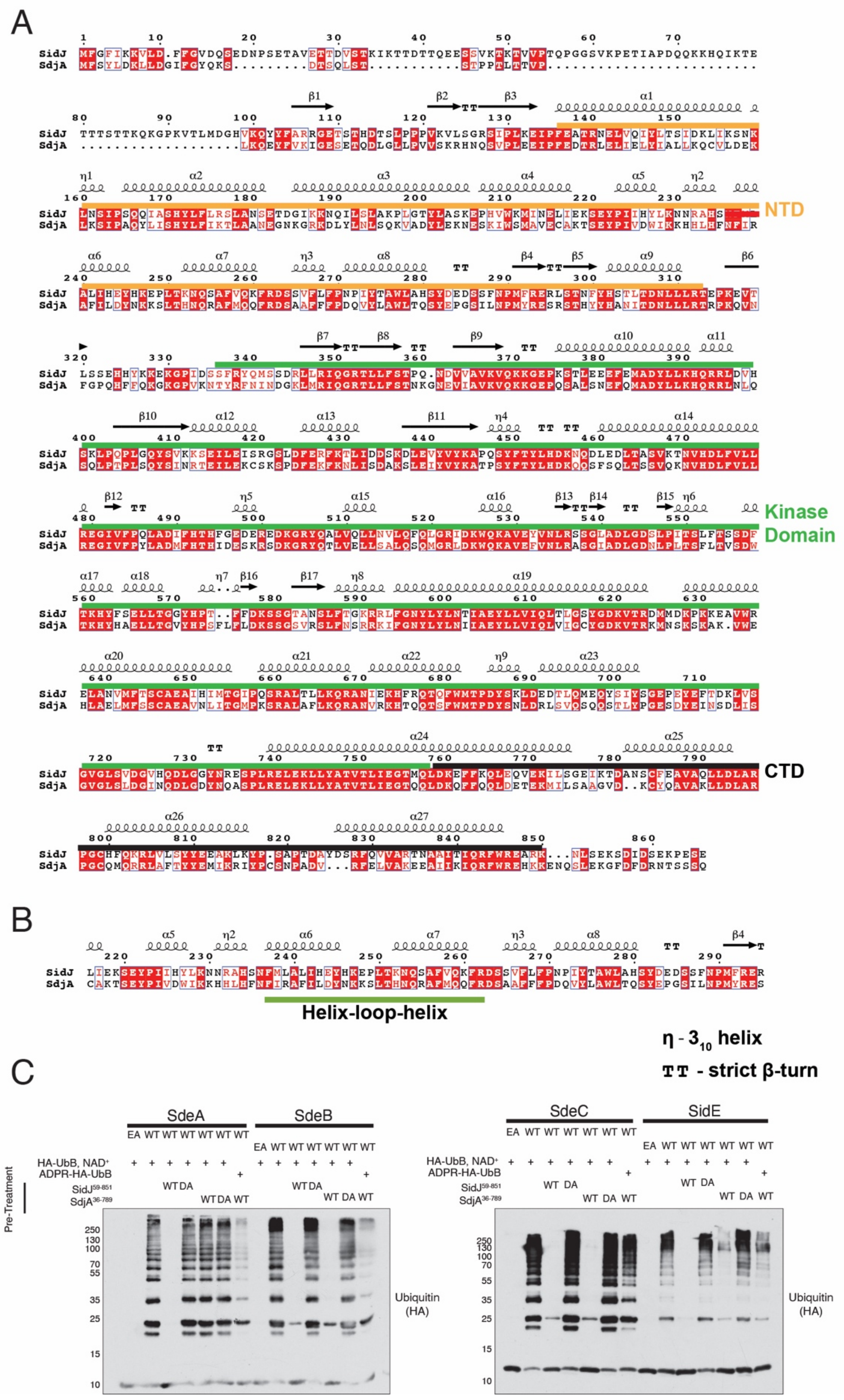
A
SidJ
SdjA
NTD
Kinase Domain
CTD
B
Helix-loop-helix
η - 3₁₀ helix
TT - strict β-turn
C
SdeA
SdeB
SdeC
SidE
HA-UbB, NAD+
ADPR-HA-UbB
SidJ59-851
SdjA36-789
Pre-Treatment
Ubiquitin (HA)

**Figure S7**. **SdjA and SidJ differentially inactivate the SidE ligases**

**(A)** SidJ and SdjA sequence comparison. SidJ and SdjA primary sequence contains an NTD, a kinase-like domain, and a CaM binding CTD. **B.** Sequence alignment with secondary structure annotated from PDBID: 6OQQ. Domain boundaries are indicated by colors: NTD (orange), kinase-like (green), CTD (black). **C.** Sequence of the helix-turn-helix motif.

**(B)** Protein immunoblotting after in vitro ubiquitination assays with the indicated mutants of SdeA, SdeB, SdeC and SidE. The SidE effectors and mutants were pretreated in glutamylation reactions with CaM in the absence or presence of SidJ$^{59\text{-}851}$/ SidJ$^{59\text{-}851;\ D542A/D545A}$ (DA) or SdjA$^{36\text{-}789}$ SdjA$^{36\text{-}789;\ D480A}$ (DA). Ubiquitination reactions were started by the addition of NAD+ and HA-Ub or HA-ADPR Ub. The reaction components were resolved by SDS-PAGE and Ub was detected by immunoblotting.

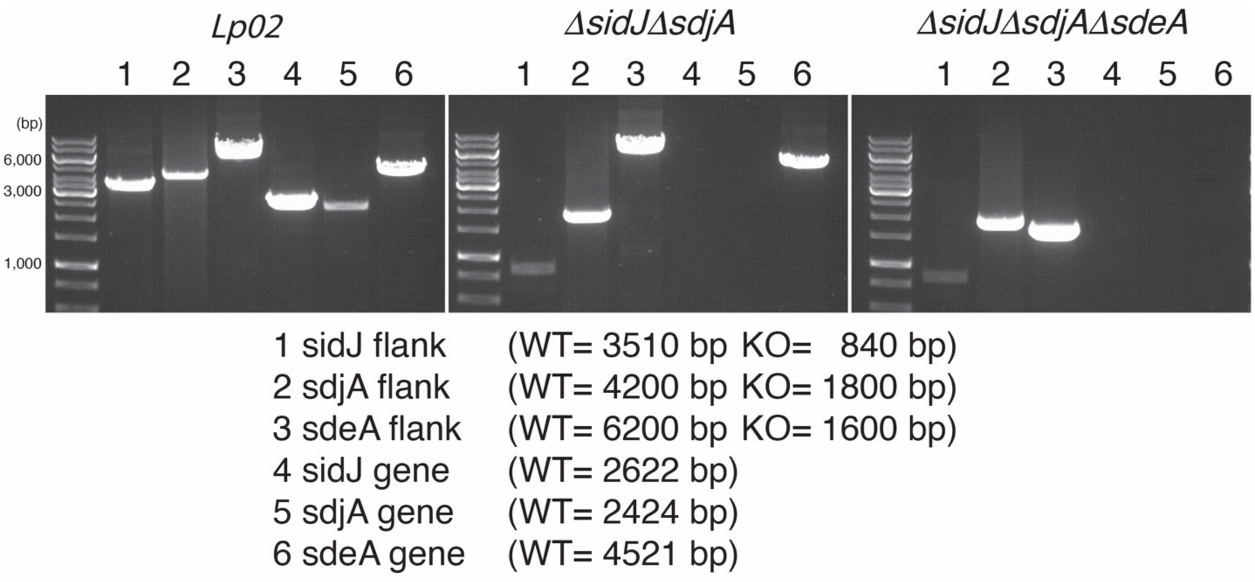


**Figure S8. Generation of *Legionella* strains.** Agarose gels depicting PCR products obtained using *Legionella* gDNA from the Lp02 (WT), *ΔsidJΔsdjA* and *ΔsidJΔsdjAΔsdeA* strains..

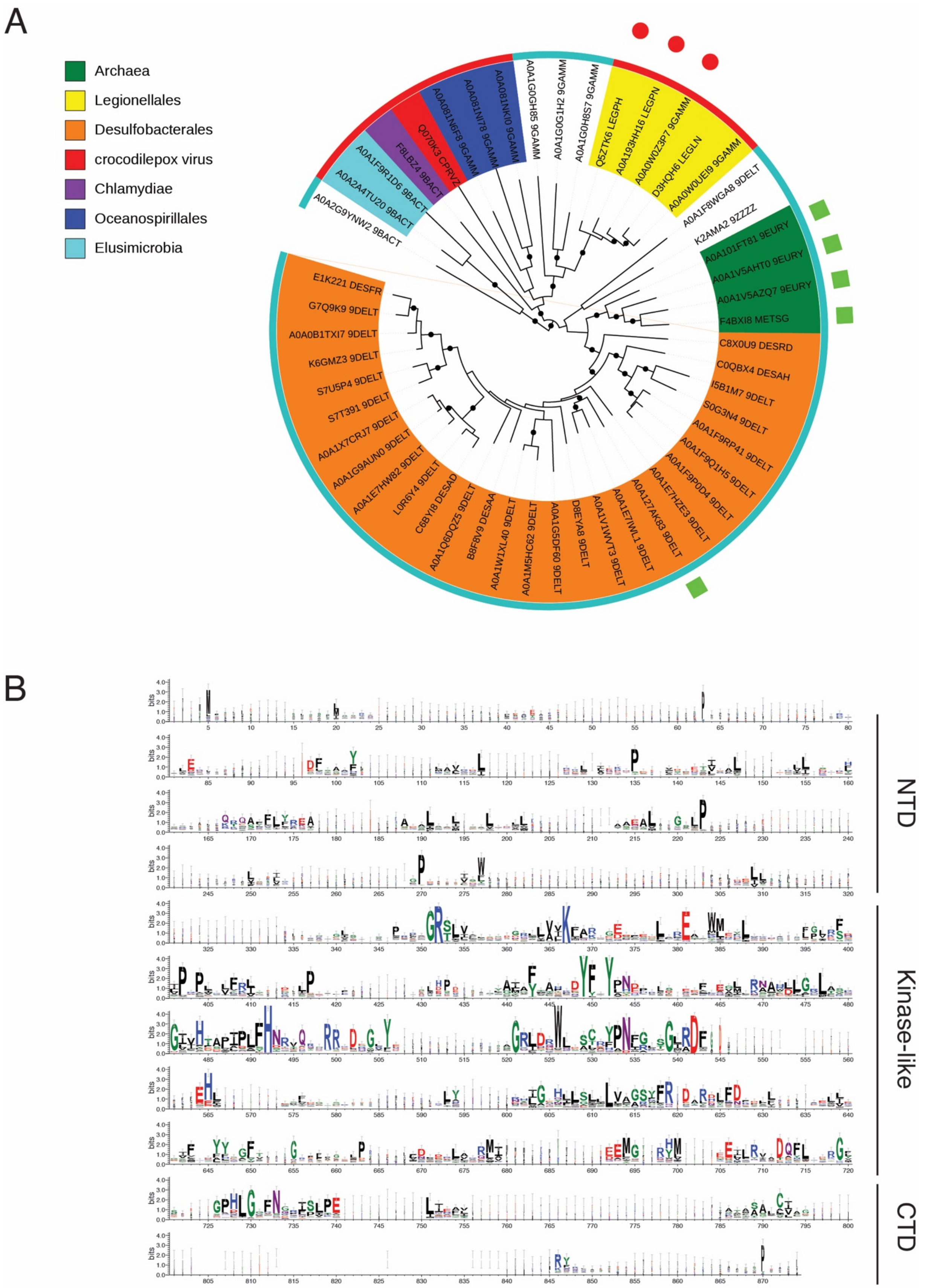
A
Archaea
Legionellales
Desulfobacterales
crocodilepox virus
Chlamydiae
Oceanospirillales
Elusimicrobia
B
NTD
Kinase-like
CTD

**Figure S9. Evolutionary analysis of SidJ.**

**(A)** SidJ-like kinase domains are found in taxonomically diverse groups of organisms. Phylogenetic tree (Maximum Likelihood) for selected SidJ homologs. Tree built from a manually corrected MAFFT multiple sequence alignment of SidJ-type kinase domains, collected using JackHMMer. Coloring by taxonomy. Red circles denote proteins containing IQ calmodulin-binding motifs, green squares denote proteins with HEAT repeats. Outer circle denotes lifestyles: red – host associated, cyan – free living, mostly aquatic. Black circles mark branches with bootstrap support above 0.9.

**(B)** SidJ-like proteins share similarity in the kinase-like domain only. Sequence logo (Weblogo) for 102 SidJ homologs collected by PSI-Blast. Logo position numbering corresponds to *L. pneumophila* SidJ. The extent of the three domains indicated. Sequences aligned by MAFFT and edited by removing columns with gaps in SidJ from *L. pneumophila*. After aligning, sequences from *Legionella* species were removed. Positions with many gaps scaled by width. Coloring according to chemical character of the residues.

**Table S1.** Data collection and refinement statistics.

| | SidJ:CaM:SdeA (EMDB) (PDB) | SidJ:CaM:SdeC (EMDB) (PDB) |
|---|---|---|
| **Data collection and processing** | | |
| Magnification | 81,000 | 81,000 |
| Voltage (kV) | 300 | 300 |
| Electron exposure ($e^-$/Å$^{-2}$) | 50 | 50 |
| Defocus range (μm) | 0.8 - 2.5 | 0.8 - 2.5 |
| Pixel size (Å) | 1.08 | 1.08 |
| Symmetry imposed | C1 | C1 |
| Initial particle images (no.) | 3,193,233 | 4,454,035 |
| Final particle images (no.) | 310,154 | 152,589 |
| Map resolution (Å) | 2.5 | 2.8 |
| FSC threshold | 0.143 | 0.143 |
| **Refinement** | | |
| Initial model used (PDB code) | 5YIM, 6S5T | SidJ:CaM:SdeA |
| Model resolution (Å) | 2.6 | 2.9 |
| FSC threshold | 0.5 | 0.5 |
| Map sharpening *B* factor (Å$^{-2}$) | -37 | -47 |
| Model composition | | |
| Nonhydrogen atoms | 13,016 | 13,039 |
| Protein residues | 1,604 | 1,616 |
| Ligands | 5 | 6 |
| B factors (Å$^{-2}$) | | |
| Protein | 49.1 | 54.9 |
| Ligands | 37.5 | 30.8 |
| R.m.s. deviations | | |
| Bond lengths (Å) | 0.002 | 0.003 |
| Bond angles (˚) | 0.438 | 0.468 |
| Validation | | |
| MolProbity score | 1.26 | 1.42 |
| Clashscore | 2.98 | 3.74 |
| Poor rotamers (%) | 0 | 0 |
| Ramachandran plot | | |
| Favored (%) | 97 | 96.2 |
| Allowed (%) | 3.0 | 3.8 |
| Disallowed (%) | 0 | 0 |

### Materials and Methods

#### Lead Contact and Materials Availability

Further information and requests for resources and reagents should be directed to and will be fulfilled by the Lead Contact, Vincent S. Tagliabracci. Plasmids, primers, recombinant protein, experimental strains and any other research reagents generated by the authors will be distributed upon request to other research investigators under a Material Transfer Agreement.

#### Antibodies

Mouse anti-GAPDH antibodies (CB1001) were from EMD Millipore. Mouse anti-HA antibodies were from Sigma (H3663) and anti-*Legionella pneumophila* antibodies (ab20943) were from Abcam. Rabbit anti-SdeA, SidJ and SdjA were raised by Cocalico and purified in-house as previously described (Sreelatha et al., 2018).

#### Generation of Plasmids and Strains

SidJ, SdjA, SdeA, SdeB, SdeC and SidE coding sequences (CDS) were amplified by PCR using *Legionella pneumophila* Philadelphia-1 strain genomic DNA (gDNA) as a template. SidJ, SdjA, SdeA, SdeC CDS were also codon-optimized for mammalian expression and synthesized as gBlocks (SidJ, SdjA, SdeC; Integrative DNA Technologies, SdeA; Genscript). CALM2 and FcγRIIa were amplified from the Ultimate™ ORF Lite human cDNA collection (Life Technologies) and cloned into mammalian or bacterial expression vectors. Amino acid mutations were introduced via Quick Change site-directed mutagenesis. Briefly, primers were designed manually, or using the Agilent Quick Change Primer design tool: https://www.genomics.agilent.com and used in PCR reactions to generate the desired mutation using PfuTurbo DNA polymerase. Reaction products were digested with Dpn1 restriction endonuclease and mutations were confirmed by Sanger sequencing. For mammalian cell expression in HEK293A cells, codon-optimized SidJ, SdjA, SdeA and SdeC were amplified from gBlocks by PCR and cloned in frame with a C-terminal V5 tag (SidJ), N-terminal V5 tag (SdjA) or amplified with an N-terminal Myc tag (SdeA, SdeC) into the CMV promoter-driven vector pcDNA

(Invitrogen). pcDNA-HA-tagged Ubiquitin B (UbB) was a generous gift from Dr. Jenna Jewell and used as a template to generate the $Ub^{GG/AA}$ mutant.

*L. pneumophila* strains Lp02, Lp03 (Lp02 ΔdotA), and thymidine auxotrophic derivatives used in this study were derived from *Legionella pneumophila* Philadelphia-1 strain and were generous gifts from Dr. Ralph Isberg. *Legionella* bacteria were maintained on ACES [N-(2-acetamido)-2- aminoethanesulfonic acid]-buffered charcoal yeast extract (CYE) agar plates or grown in ACES buffered yeast extract (AYE) liquid cultures supplemented with ferric nitrate (0.135 g/L) and cysteine (0.4 g/L). Thymidine was added to a final concentration of 100 µg/mL for maintenance of the thymidine auxotrophic strains.

SidJ, SdjA and SdeA knockout strains were generated using the R6K suicide vector pSR47s (KanR, sacB), a generous gift from Dr. Shaeri Mukherjee, UCSF. Briefly, ~800bp regions flanking the SidJ ORF were amplified and cloned using Gibson assembly into pSR47s to generate pSR47s-*ΔsdeA*, pSR47s-*ΔsidJ*, and pSR47s-*ΔsdjA*, which was maintained in S17-1 λpir *E. coli*. pSR47s vectors with flanks were introduced by electroporation into Thy- strain Lp02, or *ΔsidJ*, or *ΔsidJ/ΔsdeA*, and colonies having undergone homologous recombination were selected with kanamycin (30 µg/mL). Bacteria were streaked on CYA Thy+ plate. Metrodiploids were resolved on 5% sucrose, and the resulting colonies were screened for loss of respective genes by PCR and protein immunoblotting. Complementing strains were generated using the RSF1010 cloning vector pJB908 (AmpR tdΔi), a gift from Dr. Ralph Isberg. Mutants were cloned into pJB908 and transformed by electroporation into the appropriate parental strain. Transformants were selected for on CYE medium without thymidine and complementation was verified by PCR and protein immunoblotting. For intracellular replication assays and HEK293 cell challenge, *Legionella* were scraped from a 2/3-day heavy patch to inoculate 2 mL AYE broth. Serial dilutions of the inoculum were grown 24-48 hours at 37°C with shaking to an OD600 of 2.5 to 4, at which time the cultures acquired a brown pigmentation.

**Cell Culture**

*Acanthamoeba castellanii* was maintained as a monolayer culture in ATCC 712 PYG medium (20 g/L protease peptone, 1 g/L yeast extract, 100 mM glucose, 4 mM $Mg_2SO_4$, 0.4 mM $CaCl_2$, 0.1% (w/v) sodium citrate dihydrate, 0.05 mM $Fe(NH_4)_2(SO_4)_2 \cdot 6H_2O$, 2.5

mM $NaH_2PO_4$, 2.5 mM $K_2HPO_4$, final pH adjusted to 6.5) in tissue culture flasks at 23°C. HEK293A cells were grown in DMEM containing 10% (vol/vol) FBS with 100 μg/mL penicillin/streptomycin (GIBCO) at 37 °C with 5% $CO_2$. Cells were routinely tested for Mycoplasma contamination by PCR based methods.

### Intracellular replication in amoeba

18 hours prior to infection, confluent amoeba monolayers were washed with *A. castellani* buffer ACB (4 mM $Mg_2SO_4$, 0.4 mM $CaCl_2$, 0.1% (w/v) sodium citrate dihydrate, 0.05 mM $Fe(NH_4)_2$ $(SO_4)_2 \cdot 6H_2O$, 2.5 mM $NaH_2PO_4$, 2.5 mM $K_2HPO_4$, final pH adjusted to 6.5), to remove non-adherent cells, and collected by scraping. Resuspended in fresh ACB, amoeba were counted, and $6\times10^5$ cells were seeded into individual wells of 24-well plates 1-2 h prior to infection. Plates were equilibrated to reach 37°C. All subsequent incubations were performed at 37°C. *Legionella* cultures at post-exponential phase were diluted in *A. castellanii* buffer and ~$6\times10^4$ bacteria were added to each well for a multiplicity of infection (MOI) of 0.1 (assuming 1 $OD_{600}=10^9$ *Legionella* CFU). Infections were synchronized by centrifugation at 230 x g for 5 minutes. Infections were allowed to proceed for 1 hour, then extracellular bacteria were removed by washing each well 3 times in ACB, before adding ACB buffer to a final volume of 0.5 mL/well. Timepoint zero (1 HPI) was lysed immediately after washing, and subsequent timepoints were taken after 24h, and 48h. Amoeba cells were lysed by addition of saponin to a final concentration of 0.05%, and scraped off the plate with a mini cell scraper. Wells were subsequently washed with 500 μL of Milli-Q $H_2O$, and combined with saponin lysates. Serial dilutions of the infectious inoculum and the amoeba lysates were plated on CYE plates to confirm the MOI, quantify infection efficiency, and assess bacterial growth. All solutions used during the infection were filtered using sterile 0.22 μm syringe filters.

### Protein purification

*L. pneumophila* SidJ residues 59-851 or 97-851, SdjA FL and 36-789, human CaM (CALM2), and the SidE effectors (and indicated truncations) were cloned into a modified pET28a bacterial expression vector (ppSumo), containing an N-terminal 6X-His tag followed by the yeast Sumo (smt3) CDS. Ubiquitin, CaM, SdeA 231-1190 and SdeC 231-1222 were also cloned into pProEX2 containing an N-terminal 6X-His tag followed by a

TEV protease cleavage site. For purification from *E. coli*, plasmids were transformed into Rosetta DE3 cells. Cells were grown in Miller modification of Luria Bertani (Miller LB) broth with 100 mg/L Ampicillin or 50 mg/L Kanamycin, in presence of 34 mg/L of Chloramphenicol. At the $OD_{600}$ of 0.6-1.0, protein expression was induced with 0.4 mM IPTG for 16-18 hours at 18 °C. Cells were harvested by centrifugation and lysed in 50 mM Tris-HCl pH 8, 300 mM NaCl, 15 mM Imidazole, 1 mM PMSF, (containing 5 mM β-ME) by sonication. Cell lysates were cleared by centrifugation at 35,000 x g for 30 minutes. The cleared lysate was incubated with washed Ni-NTA beads for a minimum of one hour at 4°C. Beads were passed over a gravity column and washed with 20 column volumes of 50 mM Tris-HCl pH 8, 300 mM NaCl, 25 mM imidazole, 5 mM β-ME. Proteins were eluted with 50 mM Tris-HCl pH 8, 300 mM NaCl, 300 mM imidazole, 1 mM DTT. SdjA purifications were performed in 500 mM NaCl to aid stability. Full-length SdjA cloned into ppSumo was coexpressed with human 6X-His-CALM2 in pProEX2 and grown under triple selection (Kan, Amp, Cm), and was purified in one step (Ni-NTA). Proteins were cut overnight at 4 °C with 6X-His tagged Ulp1 Sumo protease or TEV protease, followed by gel filtration chromatography using a Superdex 200 gel filtration column attached to an AKTA Pure FPLC chromatography system (GE Healthcare) in 25 mM Tris 7.5 or 8.0, 150 mM or 300 mM of NaCl, 1 mM DTT. Proteins were concentrated (~15-30 mg/mL SidJ, ~30 mg/mL CALM2, ~30 mg/mL $SdeA^{Core}$, 60 mg/mL $SdeC^{Core}$, ~10 mg/mL full-length SidE effectors), aliquoted, flash frozen in liquid $N_2$ and stored at -80 °C until use.

**Size-exclusion chromatography for SidJ:CaM:SdeC and SdeA complexes**

For preparation of the reaction intermediate complex sample, tag-less proteins were preincubated for 15 minutes at 37 °C: 2.5 mg SidJ 97-851, 0.83 mg CALM2 and 2.5 mg $SdeA/C^{Core}$. Conditions within the preincubation were: 50 mM Tris 7.5, 50-75 mM NaCl, 10 mM $MgCl_2$, 4 mM ATP, 1 mM DTT, 300 µL. The reaction was spun at 22,000 x g for 10 minutes at 4 °C, diluted to 400 µL with size-exclusion buffer (25 mM Bis-tris pH 6.5, 100 mM NaCl, 1 mM TCEP, 2 mM $MgCl_2$, 1 mM ATP), and subjected to SEC on a Superdex S200 increase 10/300 column.

For SEC experiments, the same conditions were used, with less protein: 0.5 mg of each SidJ and SdeC and + 0.17 mg CALM2. ATP was omitted from the SEC buffer.

Preincubation was supplemented with 5 mM L-Glu for experiment with Glu treatment. Tris pH 7.5 was used for basic condition size-exclusion experiment shown in **Figure 1C**.

#### Cryo-EM grid preparation

The reaction intermediate complex sample was prepared as above, in size-exclusion buffer. All proteins were tag-less. Concentration of proteins within the complex fractions were assessed according to Bradford reagent (Bio-rad protein assay), and were ~1-1.5 mg/mL. The peak fraction was diluted to 0.35 mg/mL using size-exclusion buffer, and spun at 22,000 x g for 5-10 minutes at 4 °C. The sample used for grid preparation was never frozen and used within 1 hour after preparation. Grids were prepared using a FEI Vitrobot mark IV (ThermoFisher Scientific) at 4 °C with 95% relative humidity and plunged into liquid ethane. 3.5 µL of sample was applied to a Quantifoil 300-mesh R1.2/1.3 grid (Quantifoil, EMS), which was glow discharged using Pelco EasiGlo. The sample was preincubated with grid for 10 seconds before blotting.

#### Cryo-EM Data collection and processing

Prior to data collection, sample grids were screened on a Talos Artica microscope, and a pilot dataset of SidJ-SdeC-CaM was collected on a Titan Krios microscope at the Cryo Electron Microscopy Facility at UT Southwestern. Datasets of both SidJ-SdeA-CaM and SidJ-SdeC-CaM complexes used for final data processing were collected at the Pacific Northwest Cryo-EM Center (PNCC) on a Titan Krios microscope operating at 300 kV, with the post-column energy filter (Gatan) and a K3 direct detection camera (Gatan), using SerialEM(Mastronarde, 2005). Movies were acquired at a pixel size of 0.54 Å in super-resolution counting mode, with an accumulated total dose of 50 e-/Å$^2$ over 78 frames. The defocus range of the images was set to be -0.8 to -2.5 µm. 11,694 and 9,409 movies were collected for SidJ-SdeA-CaM complex and SidJ-SdeC-CaM complex, respectively.

#### Image processing and 3D reconstruction

Unless described otherwise, all datasets were processed with Relion (Scheres, 2012). For the pilot dataset, the motion corrected micrographs from Relion were preprocessed with MicAssess (Li et al., 2020). Particles were picked with crYOLO (Wagner et al., 2019).

A 4.5 Å map of SidJ:CaM:SdeC was obtained from the pilot dataset. This map was low pass filtered and used as the initial model for data processing as described below. For both datasets collected at PNCC, movies were aligned and summed with MotionCor2 (Zheng et al., 2017), with a downsampled pixel size of 1.08 Å. MicAssess (Li et al., 2020) was used to preprocess all motion corrected images. The CTF parameters were calculated using Gctf (Zhang, 2016), and images with estimated CTF max resolution better than 5 Å were selected for further processing. For the SidJ:CaM:SdeA dataset, 3,193,233 particles were picked using crYOLO (Wagner et al., 2019) from 9,839 images. 2D and 3D classifications identified two subclasses, which were combined with 386,460 particles. After further 3D refinement, CTF refinement and particle polishing, an additional step of alignment-free 3D classification with eight classes was performed. 310,154 particles from three subclasses with more complete maps for SdeA were combined to calculate the final map at 2.5 Å resolution. For the SidJ:CaM:SdeC dataset, 4,454,035 particles were picked using crYOLO5 from 8,432 images. 212,768 particles were selected after 2D and 3D classifications for subsequent 3D refinement, CTF refinement and particle polishing, followed by alignment-free 3D classification. 152,589 particles from the two best classes were used to calculate the final map at 2.8 Å resolution. Map resolution values were calculated by Relion with the gold standard FSC method.

**Model building and refinement**

For initial model building, the existing crystal structure of SdeA (PDBID: 5YIM), and cryo-EM structure of SidJ:CALM2 (6S5T) were fit into experimental density of SidJ:CaM:SdeA using Chimera. Models were manually adjusted in Coot to account for main chain rearrangements, compared to the crystal structure. Fit was further enhanced by rigid fit and morph routines in Phenix real space refine. Resultant model was split into SidJ:CaM and SdeA, and fit into SidJ:CaM:SdeC map using chimera. Sequence of SdeA was mutated in coot, to match SdeC sequence. Rearranged areas were manually adjusted. AceDRG from CCP4 7.1 suite(Winn et al., 2011) was used to generate AMP-Glu link restraints. DeepEMhancer(Sanchez-Garcia et al., 2020) was used to generate post-processed maps, with the two unfiltered half maps from Relion, for further manual model building and inspection in Coot(Emsley et al., 2010). The models were then subjected to real space refinement in Phenix(Adams et al., 2010), with secondary structure restraints and non-crystallographic symmetry restraints. Model geometries were assessed with MolProbity(Chen et al., 2010) as a part of the Phenix validation tools (**Table S1**). Figures were rendered in PyMOL (The PyMOL Molecular Graphics System, Schrödinger), Coot or Chimera (Pettersen et al., 2004).

**Production of SdjA antibodies**

*L. pneumophila* SdjA 36-789 D480A was purified as above as a 6X-His-SUMO fusion and was cleaved off of Ni-NTA resin with Ulp1 protease (on-column cleavage). The protein was further purified by Superdex 200 16/600 gel filtration chromatography and used to inoculate rabbits for generation of rabbit anti-SdjA anti-serum (Cocalico Biologicals). Total IgG was partially purified by ammonium sulfate precipitation as described (Kent, 1999) and affinity-purified by passing through a HiTrap NHS-activated HP column crosslinked with WT untagged SdjA 36-789. Anti-SdjA antibodies were eluted with 100 mM Glycine pH 2.5, neutralized, concentrated to ~1 mg/mL, aliquoted and stored at -20 °C until use.

**Glutamylation assays**

Glutamylation reactions were performed in a reaction mixture containing 50 mM Tris-HCl pH 7.5, 50 or 75 mM NaCl, 1 mM DTT, 1 mM $MgCl_2$, 0.3 mM ATP, and 100 μM [U-$^{14}$C] or [$^3$H] L-Glu (specific radioactivity 200 cpm/pmol for $^{14}$C, 1500 cpm/pmol for $^3$H). For standard endpoint reactions, a typical 20 μL reaction contained 10 μg SidE effector, 1 ug SidJ/SdjA, and 2 μg CaM. Reactions were initiated by adding ATP/$Mg^{2+}$ , Glu and CaM, and were allowed to proceed at 37 °C for 60 minutes, after which they were terminated by addition of 7 uL of STOP mix (0.5 M EDTA, 5X SDS-PAGE loading dye; 2:5 ratio). Reaction products were resolved by SDS-PAGE and visualized by Coomassie staining. Gels were then soaked in EN3HANCE reagent for 30 minutes, followed by a 30-minute soak in $H_2O$, then dried. $^{14}$C incorporation was detected by autoradiography.

For kinetic assays, 0.1 ug of SidJ was incubated with 10 ug SdeA$^{FL}$ for 15 minutes at room temperature (For SdjA, 0.6 ug of protein was incubated for 20 min with 5 ug of SdeC$^{Core}$). 100 μM [$^3$H] L-Glu (specific activity of 1500 cpm/pmol). Triplicate 20 uL reactions were incubated for 15 min at room temperature and terminated as above. After SDS-PAGE separation, Coomassie stained SidE bands were excised, placed in glass scintillation vials and digested in 1 mL of 30% $H_2O_2$ at 60 - 75 °C overnight. After digestion was complete, vials were cooled down to ambient temperature, mixed with 12 mL of Budget-Solve scintillation cocktail (RPI, 111167) and $^{3}$H incorporation quantified by scintillation counting.

**Adenylation assays**

SdeA adenylation end-point assays using [$\alpha^{32}$P]ATP were performed in a 20 uL reaction volume for 1 h at room temperature using 15 µg SidJ, 15 µg SdeA and 4.5 µg CALM2. Proteins were prediluted in dilution buffer (50 mM Bis-tris 6.5, 50 mM NaCl, 1 mM DTT). Reactions were initiated with ATP/$Mg^{2+}$. The final reaction conditions were: 50 mM Bis-tris pH 6.5, 50 mM NaCl, 1 mM DTT, 150 µM [$\alpha^{32}$P]ATP (SA: 1000 cpm/pmol), 1 mM $MgCl_2$, 1 mg/mL fatty acid-free BSA, -/+ 1 mM Glu. Reactions were stopped by addition of 500 µL of 30 mM ATP in 20% Trichloroacetic acid (TCA). Reactions were incubated on ice for 40 minutes, spun at 20,000 x g for 10 minutes at 4 °C and washed twice with 250 µL of 20% TCA solution. Radioactivity in the pellets was quantified by scintillation counting.

**Differential Scanning Fluorimetry/ thermal shift assay**

Protein thermal stability in solution was monitored by measurement of fluorescence of SYPRO Orange dye (Thermo). Triplicate 25 µL reactions contained 5.5 µM $SidJ^{59-851}$ and 5.5 µM CALM2. Reaction conditions were: 10 mM Tris 7.5, 50 mM NaCl, 5 mM $MgCl_2$, 1 mM ATP and 625x dilution of SYPRO Orange dye (Invitrogen, S6650). Reactions were performed in domed PCR tubes in a BioRad CFX96 thermocycler. Samples were subjected to a gradient of temperature 25-85 °C at a rate of +1 °C/minute. Fluorescent signal was read using FRET mode of the thermocycler. Melting curves were normalized within each replicate, averaged and plotted in Prism 9. The temperature at the derivative peak as reported by BioRad CFX manager, was used as the melting temperature.

**SidE Ubiquitination Assay in HEK293A Cells**

HEK293A cells were collected and plated into individual wells of a 24-well dish at near confluency. The following day, individual wells were transfected with pcDNA-HA-Ub, pcDNA-HA-$Ub^{GG/AA}$, pcDNA-SidJ-V5 (or mutants), pcDNA-Myc-SdeA -SdeB, -SdeC, or -SidE (or mutants), or empty vector using PolyJet transfection reagent (SignaGen) with a total of ~1 μg plasmid DNA per well: HA-Ub (0.5 μg): SidJ-V5 or V5-SdjA (0.5μg): myc-SdeA (0.02 μg), myc-SdeB/C/SidE (0.04 μg). DNA was premixed separately for each well and diluted with 25 μL of serum free (S.F.), antibiotic free DMEM. 25 μL of PolyJet solution in S.F. DMEM containing ~3.1 μL of PolyJet transfection reagent per well was mixed with DNA solutions and added dropwise to cells. The medium was replaced 5 hours after transfection with DMEM containing 10% FBS and Pen/Strep. ~16-20 hrs after transfection, cells were washed twice with ice cold PBS and lysed directly on the plate with 2X SDS-PAGE loading buffer (25 mM Tris-$PO_4$ pH 6.8, 20% (w/v) glycerol, 2.5% (w/v) SDS, 0.04% (w/v) bromophenol blue and 2% β-mercaptoethanol (β-ME). Samples were boiled, resolved by SDS-PAGE, transferred to a nitrocellulose membrane, and immunoblotted with the indicated antibodies.

**SidE effector in-vitro Ubiquitin laddering and inactivation assays**

Full-length SdeA, SdeB, SdeC and SidE were glutamylated in 20μL reactions containing 0.2 mg/mL SidE effector, 0.2 mg/mL, SidJ 59-851 (WT or D542A, D545A) or SdjA 36-789 (WT or D480A) and 0.1 mg/mL CALM2 in 50 mM Tris HCl pH 7.5, 50 mM NaCl, 1 mM DTT, containing 1 mM glutamic acid. Some reactions contained no SidJ/SdjA or contained catalytic mutants of each SidE effector to inactive ART activity ($SdeA^{E860Q}$, $SdeB^{E857Q}$, $SdeC^{E857Q}$ and $SidE^{E855Q}$). Reactions were initiated by addition of ATP to 1 mM and $MgCl_2$ to 5 mM and incubated for 60 min at 37°C. Reactions were then diluted 1:10 (for a final concentration of 0.02 mg/mL SidE effector) into fresh tubes containing 50 mM Tris HCl pH 7.5, 150 mM NaCl, 1 mM DTT, 5 mM EDTA and 0.15 mg/mL HA-UbB and 100 μM NAD+ or 0.15 mg/mL ADPR-HA-UbB. The phosphoribosyl-ubiquitination assay was conducted at 37°C for 10 min (for SdeA, SdeB, and SdeC) or 15 min (for SidE). Reactions were terminated by addition of SDS loading buffer with β-ME and boiling.

**Bioinformatics**

For exploration of sequence diversity among SidJ homologs, a homolog set was collected using five iterations of PSI-Blast on the NR database. In the resulting sequence set, redundancy was removed at 70% sequence identity threshold using CD-hit (Huang et al., 2010). The sequences were aligned with Mafft (Katoh et al., 2019), and in-house script removed columns with gaps in the original SidJ (*L. pneumophila*) sequence. The sequence logos were built using the Weblogo server (Crooks et al., 2004). A phylogenetic tree was built for selected SidJ-like kinase domains, using the PhyML Maximum Likelihood method (Dereeper et al., 2008) with an Approximate Likelihood Ratio Test for branch support, and visualized with iToL (Letunic and Bork, 2016).
